# Single-Cell RNA Sequencing Characterizes the Molecular Heterogeneity of the Larval Zebrafish Optic Tectum

**DOI:** 10.1101/2021.11.05.467443

**Authors:** Annalie Martin, Anne Babbitt, Allison G. Pickens, Brett E. Pickett, Jonathon T. Hill, Arminda Suli

## Abstract

The optic tectum (OT) is a multilaminated midbrain structure that acts as the primary retinorecipient in the zebrafish brain. Homologous to the mammalian superior colliculus, the OT is responsible for the reception and integration of stimuli, followed by elicitation of salient behavioral responses. While the OT has been the focus of functional experiments for decades, less is known concerning specific cell types, microcircuitry, and their individual functions within the OT. Recent efforts have contributed substantially to the knowledge of tectal cell types; however, a comprehensive cell catalog is incomplete. Here we contribute to this growing effort by applying single-cell RNA-sequencing (scRNA-seq) to characterize the transcriptomic profiles of tectal cells labeled by the transgenic enhancer trap line *y304Et*(cfos:Gal4;UAS:Kaede). We sequenced 13,320 cells, a 4X cellular coverage, and identified 25 putative OT cell populations. Within those cells, we identified several mature and developing neuronal populations, as well as non-neuronal cell types including oligodendrocytes, microglia, and radial glia. Although most mature neurons demonstrate GABAergic activity, several glutamatergic populations are present, as well as one glycinergic population. We also conducted Gene Ontology analysis to identify enriched biological processes, and computed RNA velocity to infer current and future transcriptional cell states. Finally, we conducted *in situ* hybridization to validate our bioinformatic analyses and spatially map select clusters. In conclusion, the larval zebrafish OT is a complex structure containing at least 25 transcriptionally distinct cell populations. To our knowledge, this is the first time scRNA-seq has been applied to explore the OT alone and in depth.

## INTRODUCTION

The superior colliculus (SC) is a highly laminated multisensory processing hub located in the mammalian midbrain that receives sensory input of various modalities and is involved in visual, motor, and sensory pathways ^1,2^, as well as perceptual decision-making ^3^. Despite studies dating back to the 1970s ^4^, knowledge concerning the different neuronal cell types comprising SC microcircuitries and their respective functions is limited. The homologous optic tectum (OT), found in non-mammalian vertebrate species such as zebrafish, provides an excellent opportunity to study the SC in a more physically and genetically accessible model organism ^5^. Similar to the mammalian SC, the OT has the critical role of receiving sensory input. It is responsible for visually guided behaviors such as phototaxis, prey capture, obstacle avoidance, and predator escape ^6-12^. In larval zebrafish the tectum can be broadly divided into two regions: the periventricular layer (PVL), where the majority of tectal cell bodies lie; and the neuropil (NP), consisting primarily of neurites. Within the neuropil, several layers can be further identified: the stratum fibrosum marginale (SM), stratum opticum (SO), stratum fibrosum et griseum superficiale (SFGS), stratum griseum centrale (SGC), and the stratum album centrale (SAC) ^12^. The most superficial layer, the SM, does not receive visual input, while all deeper layers are heavily innervated by retinal ganglion cells (RGCs), which project from the contralateral eye and terminate in specific tectal laminae ^13-16^. Instead, the SM receives input from the torus longitudinalis ^17,18^ while the PVL receives afferent input from the Raphe nucleus and cerebellum ^19,20^. A hypothalamic influence on the OT has also been uncovered, as neurons within the rostral hypothalamus project to the SFGS and an area between the SGC and SAC ^13^.

Although teleost fish lack a visual cortex, the tectum’s role in complex visually evoked behaviors, combined with recent evidence of an ability to respond to both auditory and water flow stimuli ^6,7,21^, attests to the functional similarities between the OT and the SC, qualifying it as a valuable, albeit simpler, model for understanding SC microcircuitry. A complete understanding of the OT at a functional and cellular level requires a comprehensive catalog of tectal neurons including their morphologies, synaptic partners and modulation, along with their molecular signatures. Such an undertaking in teleosts begun as early as the 1970s, as Meek and Schellart identified the morphologies and spatial localizations of 14 neuron types in the adult goldfish optic tectum via Golgi-stain labeling ^17^. In the zebrafish optic tectum, significant strides have been made to characterize cell type diversity using a variety of molecular and genetic methods. A Gal4 enhancer trap screen generated over 150 stable transgenic lines, 13 of which showed specific tectal expression ^22^. Three of those 13 lines were further surveyed, resulting in the identification of multiple neuronal morphologies closely resembling those previously found in the adult goldfish optic tectum ^23^. Further tectal diversity studies have resulted in the characterization of previously unidentified cell types within the optic tectum, as well as cell types that corroborate previous studies in other teleosts ^24-27^. These cell type diversity studies (which combine functional imaging, electrophysiological recordings, and neurotransmitter typing) together with the recent publication of a larval zebrafish brain atlas ^28^ constitute a wealth of knowledge surrounding the zebrafish optic tectum. However, a key piece of the comprehensive catalog is missing: the molecular transcriptomes of tectal cells. In 2020, a landmark study began this work by generating a transcriptomic single-cell atlas of zebrafish development, highlighting gene expression changes between 1 and 5 days-post-fertilization (dpf) ^29^. Undoubtedly an invaluable developmental resource, the comprehensive nature of this study precludes an in-depth analysis of the optic tectum, as it (1) looks at whole-animal gene expression changes and (2) is limited to the first few days of development. The visual system develops rapidly in zebrafish, as RGCs reach the tectal neuropil as early as 48hpf, and by 3dpf optokinetic responses are in place; however, the optic tectum undergoes rapid development between 3 and 7dpf as neurite growth and synaptogenesis continue until approximately 7-8dpf when the tectum is considered functionally mature ^30-32^. Particularly as most functional and morphological studies are conducted between 5 and 10dpf, in depth information on the molecular transcriptomes in the OT, during this time is essential.

In this study, we contribute to optic tectum characterization by fulfilling the critical need for tectal cell type classification according to gene expression profiles obtained via single-cell RNA sequencing. Single-cell RNA sequencing (scRNA-seq) technology is a relatively novel method for transcriptional analysis that has been gaining favor over the last several years. A key benefit of this technology is the ability to quantify meaningful cell-to-cell gene expression variability in diverse tissues, allowing for the creation of comprehensive cell catalogues ^33, 34^. scRNA-seq provides direct access to cellular transcriptomes, and in recent years has been successfully utilized to characterize cell populations within the zebrafish lateral line, retinal ganglion cells, and habenula ^35-37^. Using the transgenic *y304Et*(cfos:Gal4;UAS:Kaede), an enhancer trap line which comprehensively labels the optic tectum ^38^, we utilized scRNA-seq technology to identify and characterize 25 putative cell populations within the larval zebrafish optic tectum. These transcriptomic data provide an important groundwork for the complete comprehensive characterization of the OT, as we describe the neurotransmitter identity, developmental state, potential synaptic partners, and gene expression profiles of these molecularly distinct cell populations. To our knowledge, this is the first time scRNA-seq technology has been applied to the zebrafish optic tectum alone and in depth.

## RESULTS

### Transcriptomic profiling of 7dpf larval zebrafish shows that the optic tectum consists of at least 25 distinct molecular populations

To obtain OT cells for scRNA-seq, we excised and collected heads from 7dpf transgenic *y304Et*(cfos:Gal4;UAS:Kaede) larvae, an enhancer trap line which comprehensively labels the optic tecti, habenula, epiphysis, and heart; along with very sparse labeling in the olfactory bulbs (Figure 1A). Similar to previous scRNA-seq characterizations in zebrafish ^35-37^, we found enzymatic dissociation with gentle trituration and fluorescence-activated cell sorting (FACS) at minimal flow rate to be an effective and robust method for preparing single cell suspensions for scRNA-seq (Figure 1B). To preserve transcriptional expression, we immediately methanol fixed cells post-FACS.

**Figure 1.**
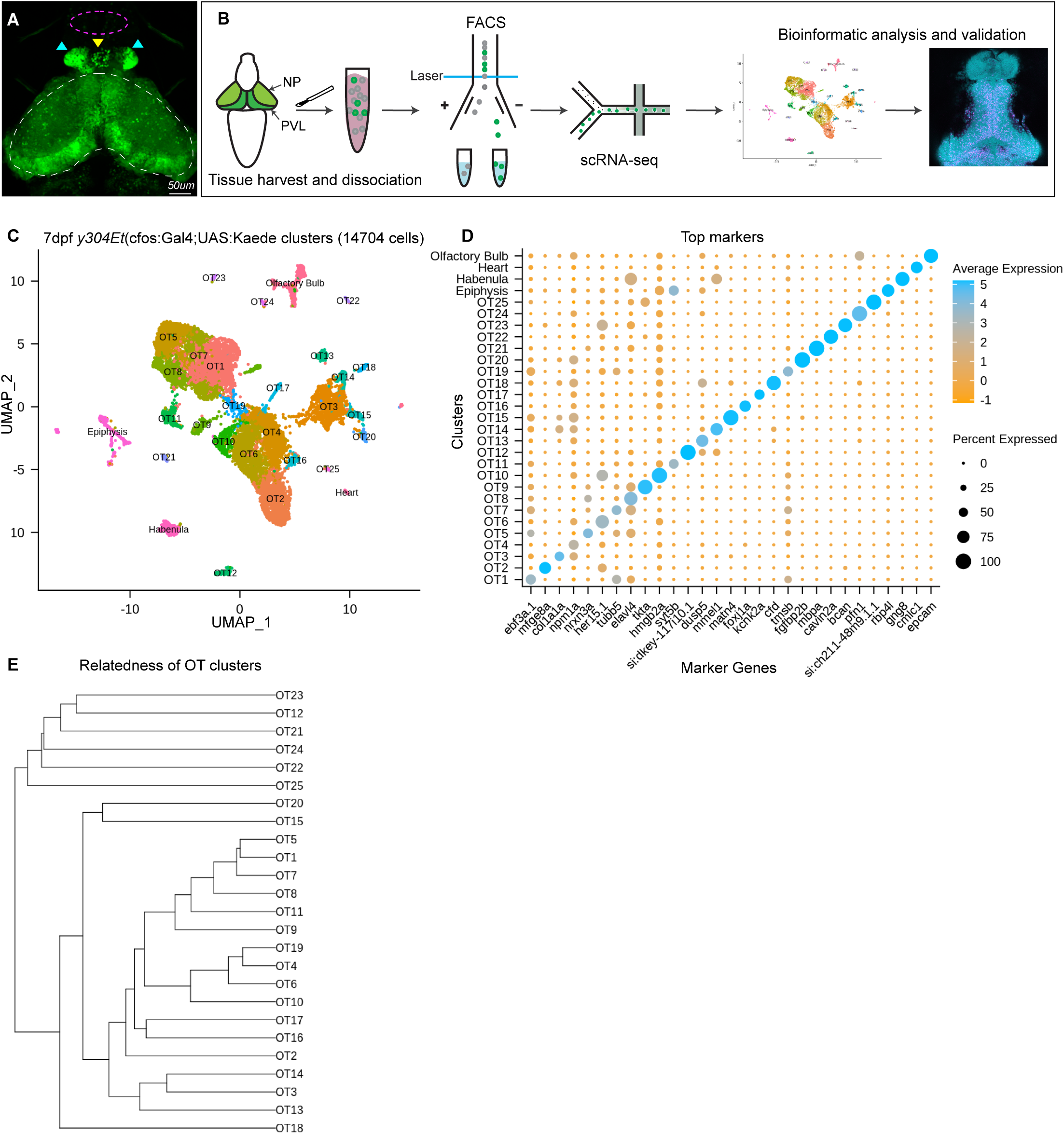
Analysis of scRNA-seq Data Reveals 25 Distinct Populations Within the Larval Zebrafish Optic Tectum. A: 7dpf representative image of the *y304Et*(cfos:Gal4;UAS:Kaede) enhancer trap line used to obtain tectal cells. Optic tectum (white dashes), habenula (cyan arrows), epiphysis (yellow arrow), and olfactory system (magenta dashes) are indicated. The heart is not shown. B: Experimental workflow for the collection and isolation of tectal cells. Heads from 7dpf Kaede^+^ larvae were collected and enzymatically dissociated, followed by FACS to isolate fluorescent cells. After quality control, 14,704 single-cell libraries obtained by the 10X Genomics Chromium platform were used for downstream analysis. PCA and graph-based clustering were utilized to sort cells into clusters and identify candidate marker genes. Bioinformatic validation and spatial mapping was completed via fluorescent HCR RNA-FISH. C: Uniform manifold approximation and projection (UMAP) showing cell populations identified via scRNA-seq analysis of 14,704 Kaede^+^ cells. Each point represents a single cell colored according to cluster identity as determined by unbiased graph-based clustering using the Louvain algorithm with multilevel refinement. UMAP embedding was used for visualization purposes only and not to define the clusters. The resulting two-dimensional spatial arrangement of clusters is a product of genetic similarity, as similar clusters are represented as closer together. D: Candidate marker genes (columns) for each cluster (rows) were nominated by differential expression analysis wherein candidate genes must: (1) be expressed in at least 50% of cells within that cluster, and (2) show specificity to the cluster compared to all other cells by exhibiting ≥1 log2 fold-change in expression with at least 25% difference in gene presence. E: Phylogram showing cluster relatedness based upon variable features defined by SCTransform and calculated within PCA space.

Using the droplet-based 10X Genomics Chromium platform, we sequenced RNA from 15,922 cells with an average of 1,419 transcripts and 648 genes recovered per cell. We filtered out low-quality cells by removing those with fewer than 500 UMI tags and 300 genes per cell, as well as cells with greater than 10% mitochondrial content. Following quality control, we scaled and normalized 14,704 cells and identified 33 distinct populations. Through differential expression analysis we nominated sets of candidate marker genes that show specificity to their respective cluster. We used known marker genes for the heart, habenula, epiphysis, and olfactory system (Table S1) to remove these populations from downstream analysis, and designated the remaining 25 populations (13,320 cells) as putative tectal cells. Although many clusters could be uniquely identified with a single marker gene (Figure 1D), in several instances clusters were best described using the top 5-10 differentially expressed genes (DEGs) (Table S2). The identification of clusters in scRNA-seq data is typically accompanied by manual annotation of known cell types. However, prior to this study the transcriptomic knowledge of tectal cells has been limited, disallowing this method of cluster validation. To preclude over-clustering based on technical noise, we identified clusters based upon the presence of marker genes meeting our diagnostic criteria and concluded that at 7dpf the larval zebrafish brain contains at least 25 transcriptionally distinct cell populations. Due to this conservative approach, it is possible that several OT clusters may include multiple cell types.

### Fluorescent in situ hybridization validates bioinformatic findings and spatially maps populations of interest

To validate our transcriptomic and clustering data, we performed hybridization chain reaction RNA fluorescence *in situ* hybridization (HCR RNA-FISH) using custom probes for *gng8, nrxn3a*, and *robo4* (Molecular Instruments, Los Angeles, CA). *gng8*, a well-known habenula marker, was used among other determinant genes to annotate and remove habenular populations (SI Figure 5A). HCR RNA-FISH targeting of *gng8* confirmed that its expression is limited to the habenula (Figure 2A-A”), validating the clustering algorithm (Figure 2B) and our method of cell exclusion. In addition to bioinformatic validation, we used HCR RNA-FISH labeling to spatially map previously uncharacterized populations. HCR RNA-FISH of OT5 marker *nrxn3a*, (Figure 2C-C”, 1D), confirmed expression is found in the OT. Moreover, it showed that *nrxn3a*^*+*^ cells are primarily found deep within the periventricular layer, with most located near the intratectal commissure that separates each tecti (Figure 2C’). Additionally, it is important to note that although *nrxn3a* is expressed higher within OT5 than other populations, it is not exclusive to it (Figure 1D); thus, we anticipate HCR RNA-FISH labeling includes a portion of OT8 cells as well. Lastly, we performed HCR RNA-FISH of *robo4*, a marker that labels 51% of the OT2 cluster (Table S2, Figure 2F), and found its expression to be limited to cells in the tectal proliferation zone (Figure 2E-E’’). Taken together, HCR RNA-FISH validated our bioinformatic method of excluding cells of non-interest and shows its capability for high resolution labeling and spatial mapping of OT populations of interest.

**Figure 2:**
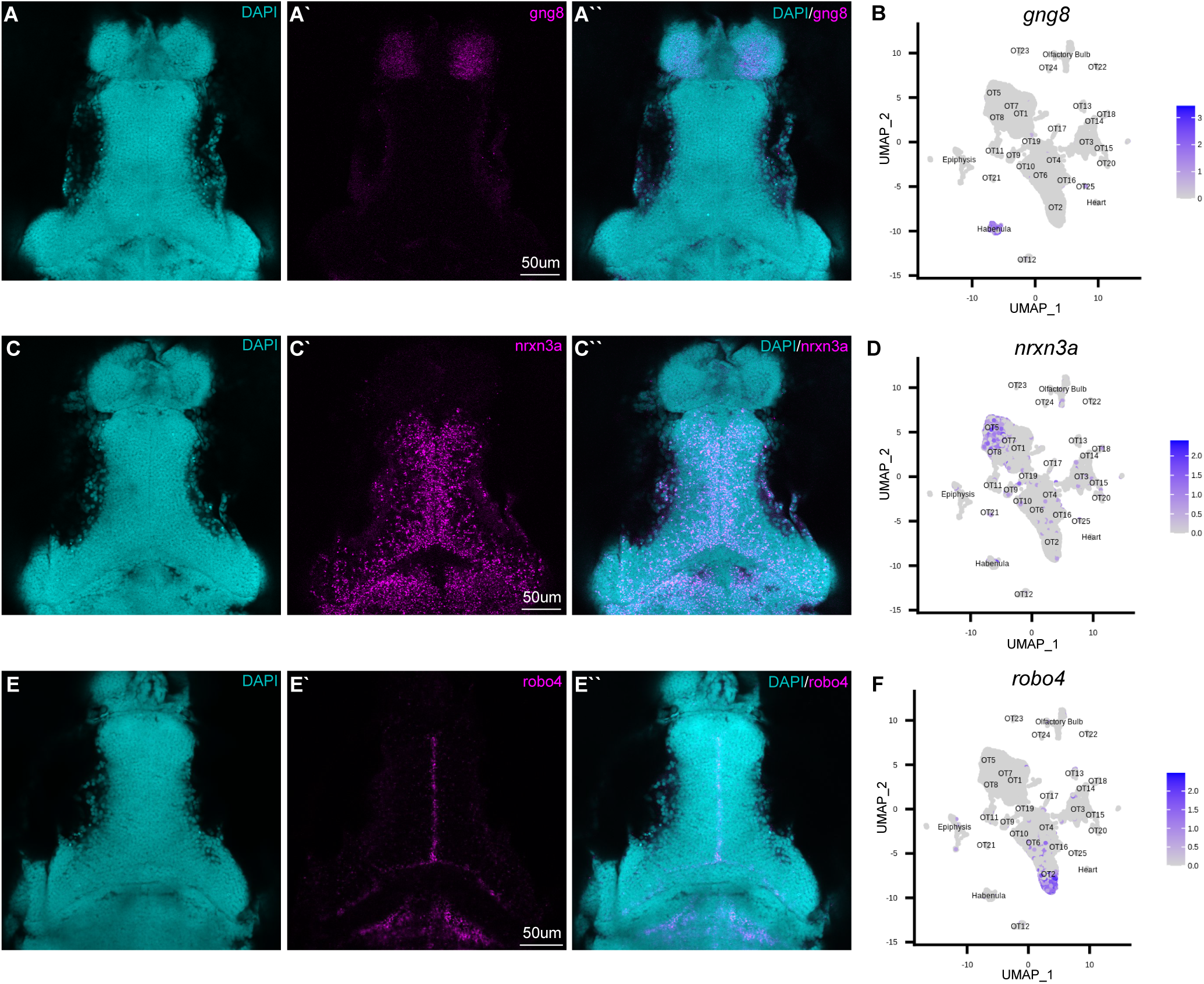
HCR RNA-FISH validates bioinformatic findings and spatially locates tectal populations. Hybridization chain reaction RNA-fluorescence *in situ* hybridization (HCR RNA-FISH) was performed using proprietary custom probe sets designed by Molecular Instruments to spatially map populations and validate bioinformatic exclusion of non-interest cells. A-A’’; C-C’’, E-E’’: Single z-slices of representative images showing expression of habenula (*gng8*) and OT (*nrxn3a, robo4*) markers in 7dpf larval zebrafish. A-A’’: *gng8*^+^ cells are located in the habenula, validating *gng8* expression as a method for annotating habenular cells and spatially mapping 89% of habenula cells (SI Table 2). B: Feature plot showing *gng8* expression is restricted to the annotated habenula cluster. C-C’’: *nrxn3a*^+^ cells are primarily located medially within the periventricular layer, and spatially map 54% of OT5 (SI Table 2). D: Feature plot showing expression of *nrxn3a* is enriched in OT5 and to a lesser extent, OT8. E-E’’: *robo4*^+^ OT cells are located near the intratectal commissure, likely within the tectal proliferation zone, and spatially map 52% of OT2 cells (SI Table 2). F: Feature plot showing *robo4* expression is restricted to OT2. All HCR RNA-FISH panels include DAPI staining as a cellular reference.

### The larval OT contains both mature and developing neurons as well as several glial populations

As a preliminary survey, we performed whole-dataset Gene Ontology (GO) analysis to identify which biological processes are upregulated in the larval optic tectum at 7dpf. Although GO analysis is often used to explore gene expression differences between conditions; it can be a valuable exploratory tool when approaching characterization of an unknown structure or developmental stage. We implemented the MAST test to identify 2,222 DEGs that were used as input for GO analysis (Table S2) and found upregulated processes such as *myelination, synapse organization, axon guidance*, and *synaptic vesicle endocytosis* (Figure 3A, Table S3), suggesting the presence of mature neurons and glial cells, as well as several immature or developing populations. However, whole-dataset GO analysis cannot show which populations are contributing to particular processes, and due to large discrepancies in size, individual cluster analysis can misrepresent data during the overrepresentation comparison. For that reason, we used GO analysis purely as an initial screening tool to guide further gene expression explorations. We used marker genes for mature neurons (Table S4) to identify OT1, OT8, and OT9 (among others) as mature post-mitotic neuronal populations (Figure 3D). Several clusters, notably OT2, OT5, and OT10, also show upregulated neurogenesis genes (Figure S5), suggesting that at this stage the OT is still developing.

**Figure 3.**
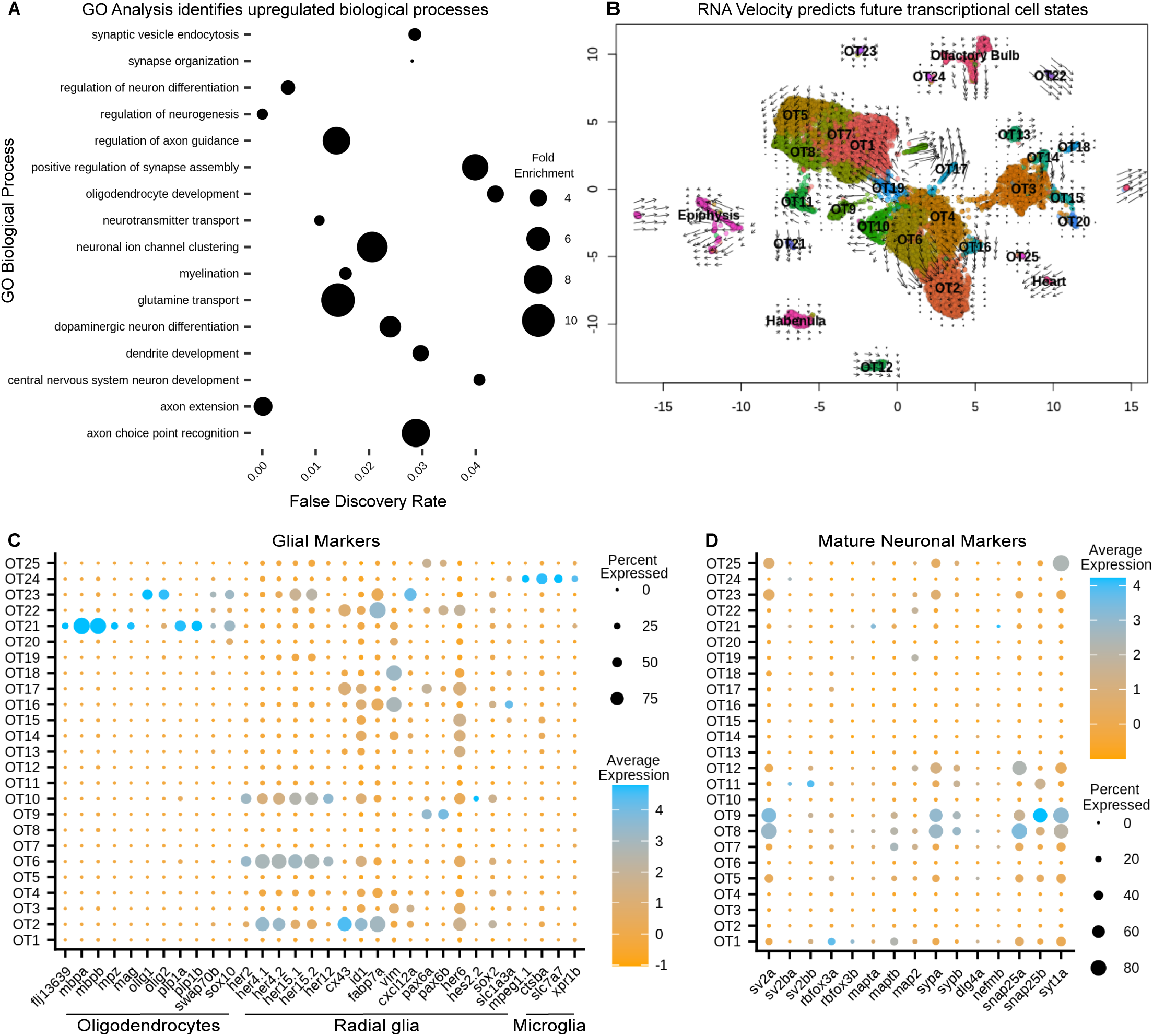
Gene Ontology (GO) and RNA Velocity Analysis Reveals Upregulated Biological Processes and predict future transcriptional cell states. A: Upregulated processes in all 13,320 putative tectal cells. Whole dataset GO analysis was performed on all significant (p<0.05) differentially expressed genes (See also Table S2). Processes of note include regulation of axon guidance, positive regulation of synapse assembly, glutamine transport, and oligodendrocyte development, suggesting the presence of immature/mature neuronal populations and glial cells (See also Table S3, S6). B: RNA velocity with original UMAP cluster embeddings. Shorter arrows or dots connote more mature cells. Arrows are vectors where direction indicates the predicted transcriptional state and magnitude indicates the degree of differentiation. BAM files from two temporal replicates were used to generate a matrix containing spliced:unspliced count ratios for each gene, where higher spliced counts indicate a more processed form of the gene and lower spliced counts indicate newly “born” genes. C: Cluster analysis of glial markers identifies OT21/OT23 as oligodendrocytes; radial glial markers are found in various populations including OT6 and OT10; OT24 is likely microglial (See also Table S5). D: Cluster analysis of manually curated neuronal markers (See STAR Methods) shows several mature populations including OT8 and OT9 (see also Table S4).

In addition to mature and developing neuronal populations, we characterized the OT21 and OT23 clusters as oligodendrocyte populations, demonstrated by their enrichment for *mbpa/b* and *olig1/olig2*, respectively (Figure 3C). In addition to oligodendrocytes, we identified several populations enriched for radial glial markers, notably clusters OT2, OT6, and OT10 (Figure 3C). Lastly, we identified OT24 as microglial, due to expression of *mpeg1*.*1, slc7a7*, and *ctsba* (Figure 3C), the latter a lysosomal gene found to be enriched in tectal populations of microglia ^39^ (See Table S5 for all glial markers). Thus, at 7dpf the larval optic tectum contains at least three non-neuronal cell types and 22 neuronal populations. Of these, we consider 7 to be mature and 15 to be immature. However, neurodevelopment is a process, and some of these immature populations are further along their developmental trajectory than others. A summary of cell type classification, developmental stage, and neuronal profile (if applicable) is found in Table 1.

**Table 1:**
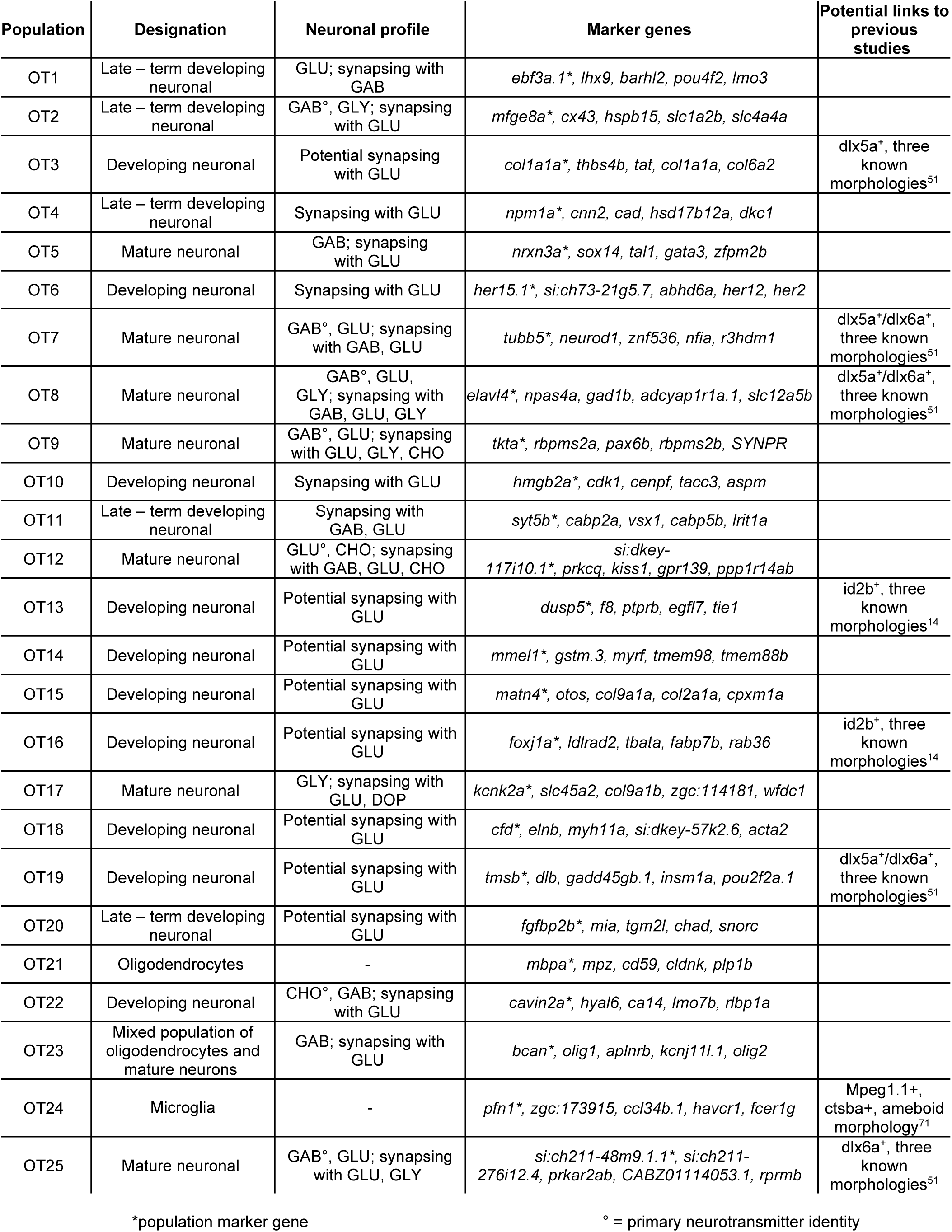
Summary of 25 transcriptionally distinct tectal cell populations. A brief overview of transcriptomic findings from targeted scRNA-seq of the larval zebrafish optic tectum. GLU = glutamatergic, GLY = glycinergic, GAB = GABAergic, CHO = cholinergic, DOP = dopaminergic. Bolded identifiers indicate the primary neurotransmitter expressed in that population

### Trajectory analysis via RNA velocity can predict future transcriptional cell states

To determine current and future transcriptional cells states of OT clusters, we performed RNA velocity analysis, which uses the ratio of unspliced to spliced gene transcripts of a gene (or proportion of intron retention), to estimate the directionality and magnitude of transcriptional changes on the timescale of hours ^40^. Conceptually, a higher ratio of unspliced:spliced indicates “newer” genes that are actively upregulated, while a lower ratio indicates mature, highly processed transcripts. This distinction is what enables RNA velocity analysis to infer current and future transcriptional cell states for each cluster. Thus, we calculated and displayed velocity estimates on our original UMAP to visualize predicted transcriptional cell states (Figure 3B). Superimposed dots indicate mature, terminal populations, while arrows can be interpreted as vectors, where the direction and magnitude indicate the predicted future transcriptional state. Notably, clusters OT8, OT24, and OT25 appear to be terminally differentiated while OT6, OT16, and OT22 are actively differentiating, evidenced by superimposed arrows over these clusters pointing to OT2 (Figure 3B). This corroborates our previous assessment that these populations are likely developing neurons and predicts that if development were to proceed normally, they may eventually transcriptionally resemble OT2. It is important to clarify that RNA velocity does not predict cell identity, and we cannot conclude that OT6 will develop into OT2. However, it may eventually share transcriptional similarities such as a GABAergic identity (Figure 4A), but all GABAergic neurons are not identical in morphology, circuitry, or function. Similarly, we do not anticipate all OT2 cells to be mature due to an upregulation of radial glial genes and neurogenesis markers (Figure 3C; SI Figure 5). Taken together, our RNA velocity results corroborate immature and mature population findings and build upon those observations by predicting the future transcriptional state of differentiating populations.

**Figure 4.**
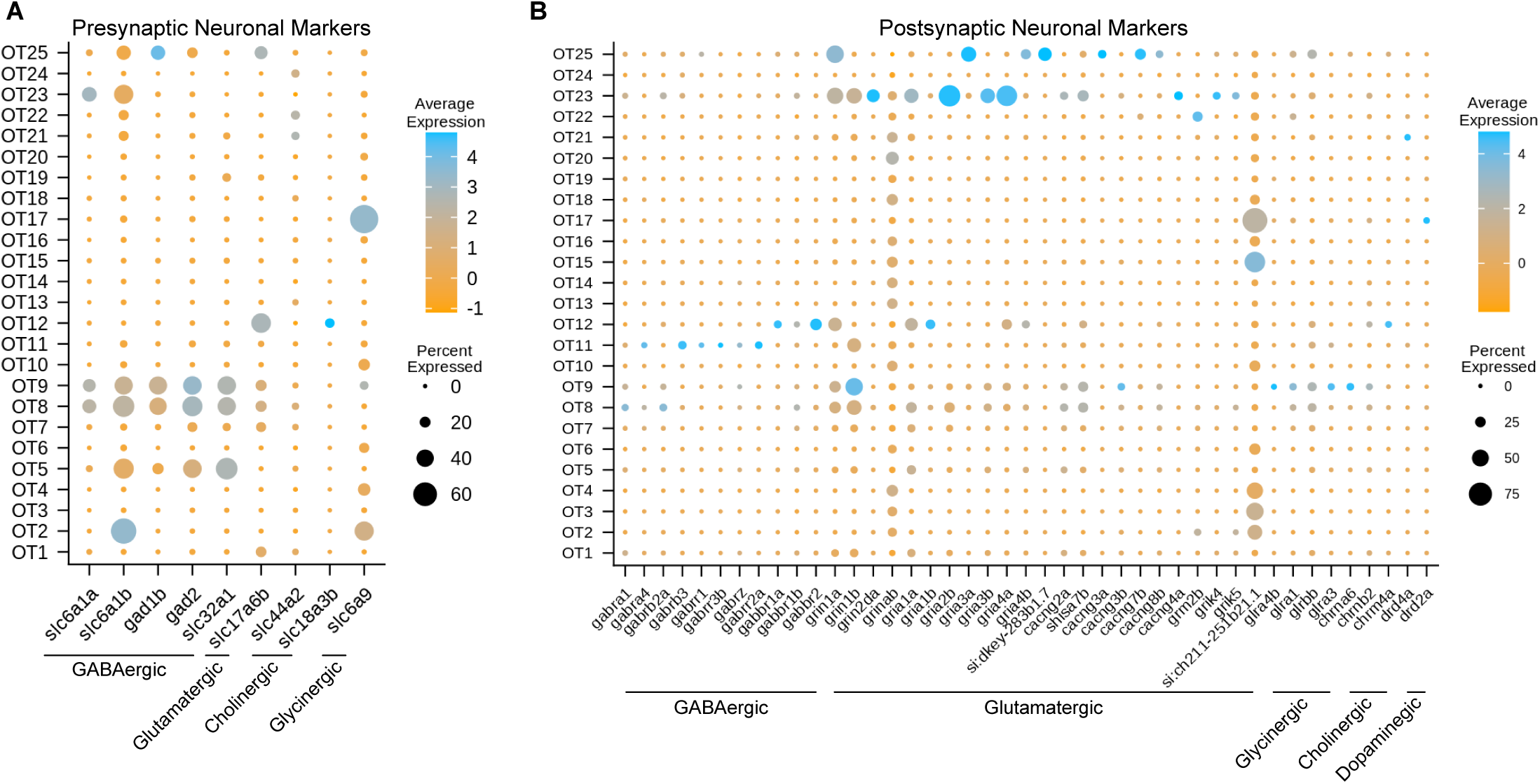
Gene Expression Profiles Characterize the Neuronal Profiles of OT Cells. A: Select presynaptic markers identify mature inhibitory (GABAergic, glycinergic, cholinergic) and excitatory (glutamatergic) tectal populations. B: Select postsynaptic markers identify the potential synaptic partners of tectal neurons by exploring genes required for various neurotransmitter receptors. All markers were manually curated based on functional designation on the ZFIN database (https://zfin.org/); see Table S7 for all pre- and postsynaptic markers.

### The majority of mature tectal neurons send inhibitory signals and receive excitatory input

To determine the neuronal profiles of mature tectal neurons, we used pre- and postsynaptic marker genes for different neurotransmitter types (SI Table 7). We determined that at 7dpf, the zebrafish optic tectum contains a strong presence of mature GABAergic neurons, and to a lesser extent, glutamatergic and glycinergic neurons (Figure 4A). This is consistent with previous studies describing GABAergic neurons as the majority population ^25,28^. While dopaminergic and serotonergic populations were not observed, GABAergic population OT9 is of particular interest, as those cells express glutamatergic, glycinergic, and cholinergic receptor-building genes (Figure 4B; Table 1); suggesting that this population receives input from various neuronal types. Interestingly, we found OT17 to highly express *slc6a9* (GLYT1), which encodes a transporter responsible for presynaptic re-uptake of the inhibitory neurotransmitter glycine. Glycinergic neurons have previously been described in the optic tectum of the lesser-spotted dogfish ^41^, and in non-neuronal cells of the hindbrain and spinal cord in zebrafish ^42^. Overall, we found major GABAergic populations such as OT8 AND OT9 to differentially express glutamatergic receptor genes, indicating their main synaptic partners are excitatory.

### OT populations differentially express transcription factors

Often overlooked in scRNA-seq analysis, transcription factors are the master regulators of gene expression and offer valuable cell identity information concerning the establishment and maintenance of cell fate. To explore these factors within OT populations, we obtained a list of 310 (257 present in data) zebrafish transcription factors from the UniProt Consortium ^43^ to determine the top overall expressed and top differentially expressed factors in each population. We identified the top two differentially expressed transcription factors (Figure 5A) within each cluster; meaning they are expressed more highly within that population as compared to all other cells, not that they are unique to that population alone. Of particular interest is the strong upregulation of *jdp2b* in OT2 (Figure 5A). *jdp2b* protein represses AP-1, a separate transcription factor that functions in cell proliferation and differentiation ^44^, indicating OT2 may be nearing its developmental end. This is consistent with previous conclusions identifying OT2 as a developmentally heterogeneous population, with indications of both developing (Figure 3C; SI Figure 5), and mature features (Figure 4A, 4B).

**Figure 5:**
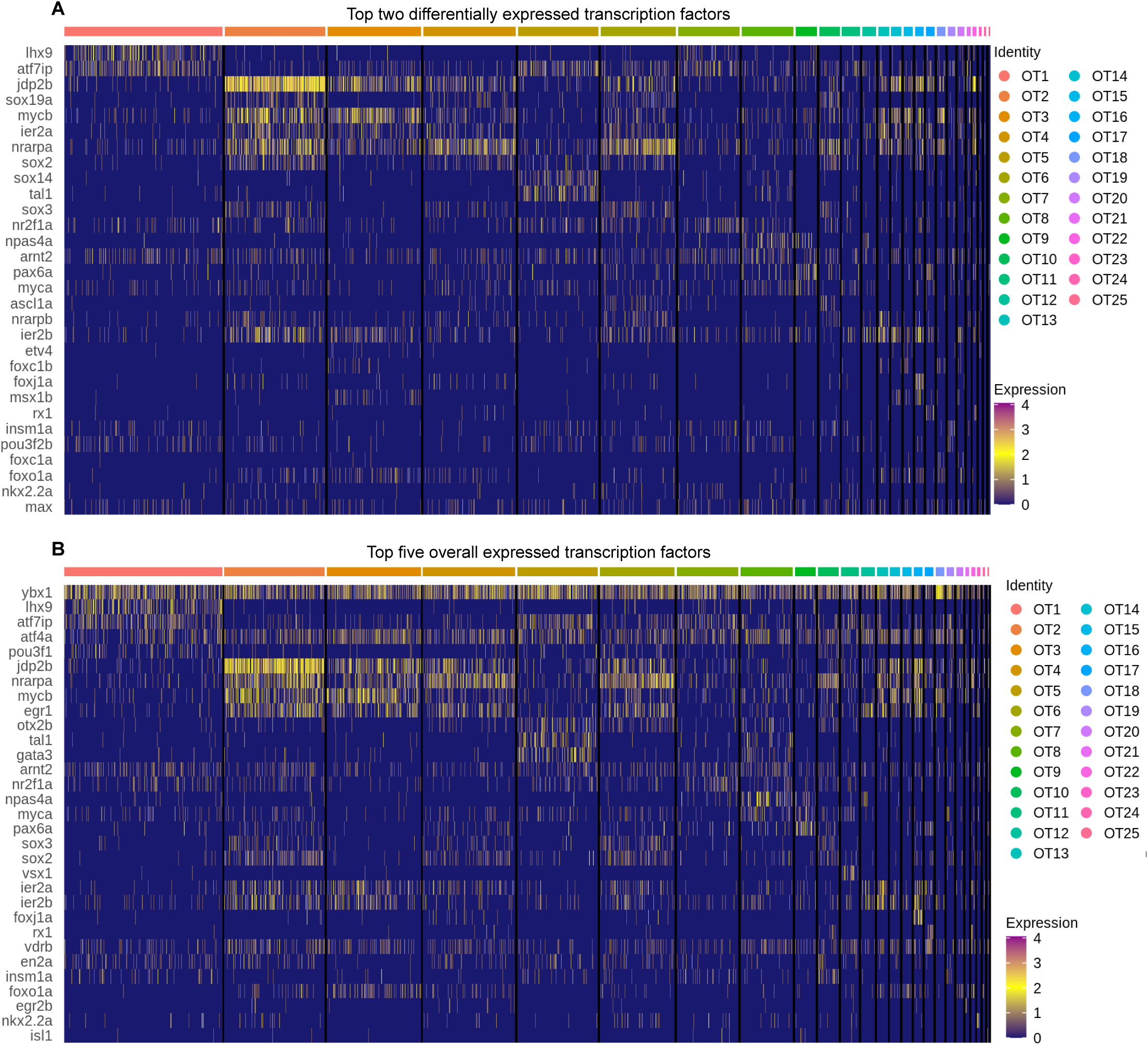
Top transcription factors in the larval zebrafish OT: The top two differentially expressed transcription factors in each OT population as compared to all other cells. B: Top five expressed transcription factors in each population. All: Duplicates of top factors are allowed but displayed only once. Factors were considered differentially expressed for a particular cluster if they were expressed >0.25 log2 fold change above all other cells and are present > 25% of cells within the cluster. Factors were ordered in decreasing expression (log2fc) level and the top two or five were selected for visualization; see Table S8 for all transcription factors.

In contrast to the relative specificity of *jdp2b* to OT2, we observed many cell populations that express the same or similar transcription factors. However, it is highly unlikely that these factors modulate gene expression in identical manners across the entire optic tectum. Rather it is the unique combination of multiple factors, coupled with developmental stage and tissue, which work together to establish and maintain that cell’s identity. To include this likelihood in our characterizations, we not only looked at differentially expressed transcription factors, but also top expressed (in terms of percentage of expressing cells) in each cluster. We found several factors, *ybx1* and *atf4a*, to be fairly ubiquitous across all OT populations (Figure 5B) despite the molecular heterogeneity of OT cells. In this regard, *ybx1* and *atf4a* pose an interesting avenue for further investigations regarding regulation of tectal cell identity.

### Tectal cell populations show an upregulation of genes implicated in neurodevelopmental disorders

The optic tectum, like its mammalian counterpart, the superior colliculus (SC), is a sensory processing hub that receives multimodal stimuli and is responsible for eliciting appropriate behavioral response ^1^. Recently, the SC has received attention for a proposed role in autism spectrum disorder (ASD) pathogenesis due the essential role it plays in sensory perception ^45^. Considering the homologous nature of the optic tectum, we investigated the tectal expression of 316 genes (407 zebrafish orthologs) implicated in ASD pathogenesis (SI Table 8). Of those, 348 were present in the OT to some degree, while 80 were considered differentially expressed (>0.25 log2fc, >25% of cluster cells). We found several genes, such as *seta* and *eif3g*, to be expressed consistently across all clusters, while others showed greater cluster specificity, such as *prex1* (OT2), *nrxn3a (*OT5), *aspm* (OT10) and *nr4a2a* (OT12) (Figure 6). This population-specific differential expression of ASD-implicated genes highlights the potential for investigations into the hypothesized link between the superior colliculus (or optic tectum) and ASD pathogenesis.

**Figure 6:**
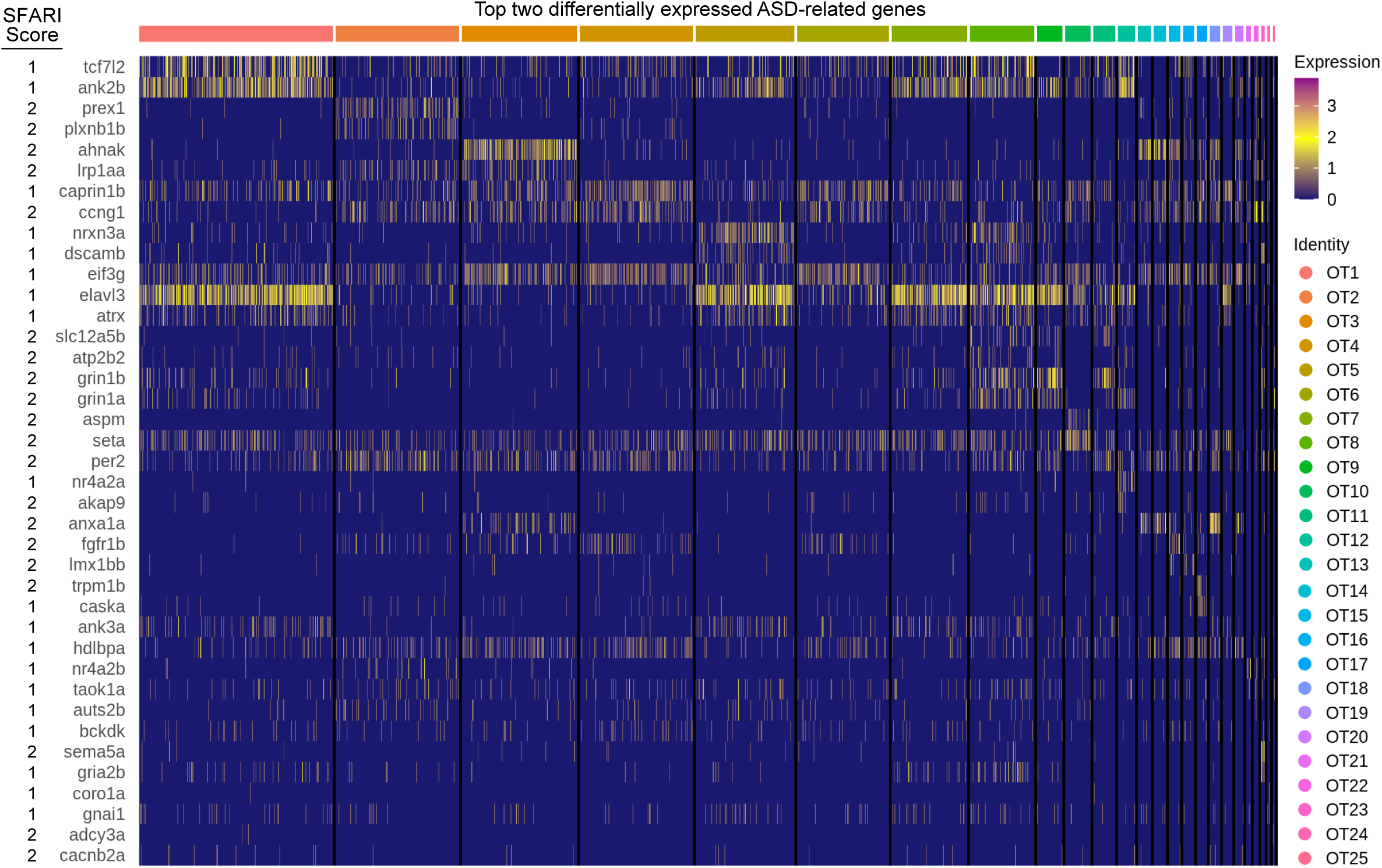
ASD implicated genes are differentially expressed in the larval zebrafish OT. The top two differentially expressed genes implicated in ASD pathogenesis, duplicates are allowed but only displayed once. Score: indicates ASD association score conferred by SFARI Gene. See Table S8 for all ASD implicated genes and their respective scores.

## DISCUSSION

To compliment current tectal characterization efforts, we describe the transcriptomic profiles for 25 molecularly distinct tectal cell populations including novel marker genes, neurotransmitter identities, potential synaptic partners, transcription factor expression, and developmental predictions.

### Developmental state of the optic tectum at 7dpf

Previous functional studies have shown escape, prey detection, and orientation behaviors are present in larval zebrafish ^9^, indicating that at 7dpf the OT has some level of functional maturity. Our data supports this through the identification of several mature neuronal populations; however, strong expression of radial glia markers across multiple clusters is notable and may indicate immature populations. As mature radial glia give rise to neurons, many markers are simply neurogenesis genes, and their expression may denote immature neurons rather than true mature radial glia. At 5dpf, the larval OT contains a proliferation zone that functions as a post-embryonic neurogenic niche that gives rise to neurons through transient radial glia cells expressing *her4* ^46^. Our data shows OT2, OT6, and OT10 express several *her4* genes including *her4*.*1* and *her4*.*2*. Additionally, HCR RNA-FISH of *robo4*, which roughly marks half of OT2 cells, spatially maps these cells to the proliferative region of the OT. In mice, *robo4* plays a role in the radial migration of later born neurons within layers II/III of the developing cortex ^47^. Localization of *robo4* in the OT’s proliferative zone may indicate it plays a similar role in the migration of post-embryonic newborn neurons within the optic tectum.

### Neuronal cell types in the larval optic tectum

Despite a wealth of data concerning tectal cell characterization, many studies are conducted using enhancer trap lines and have limited gene expression information despite excellent morphological characterization. Although we did not determine the morphologies of each population, we demonstrate the ability to connect our transcriptomic data with previous work by highlighting several highly influential studies that characterized various tectal cell types (Table 1).

In 2011, Robles et al. utilized a transgenic line labeling *dlx5a*/*dlx6a* positive neurons in the larval zebrafish optic tectum, identifying three morphological classes: GABAergic non-stratified periventricular interneurons (nsPVINs), glutamatergic bi-stratified periventricular interneurons (bsPVINs), and GABAergic periventricular projection neurons (PVPNs) ^25^. In our data, 5 populations express either *dlx5a, dlx6a*, or both. It is possible that nsPVIN and PVPN morphologies denote cells within GABAergic OT7 or OT8 clusters, which express both *dlx5a*/*dlx6a*. bsPVIN morphologies may be found in OT25, which expresses *dlx6a* and contains glutamatergic neurons. However, as many neuronal populations are still immature at 7dpf, these morphologies may also be found in populations such as OT19, which expresses both *dlx5a*/*dlx6a* but is still developing.

A second study conducted in 2019 used the *id2b:Gal4* transgene to label a subset of tectal neurons in larval zebrafish ^26^. Similar to the Robles study, genetic mosaic labeling of single neurons led to the identification of three tectal morphological types (in descending proportion): pyramidal neurons (PyrNs), projection neurons that project to the torus longitudinalis (TLPNs), and a second projection neuron type that projects to the tegmentum (TGPNs). Surprisingly, most *id2b*(*EGFP*)^+^ cells appear to be cholinergic (likely PyrNs), while approximately one-third are glutamatergic (likely TLPNs), and only 10% are GABAergic (likely TGPNs) ^26^. We found few cholinergic neurons; however, two developing populations without a known neurotransmitter identity show expression of *id2b*, thus we hypothesize these populations may have PyrN, TLPN, or TGPN morphologies.

Superficial interneurons (SINs) have also been described as primarily GABAergic and receiving glutamatergic input directly from retinal ganglion cells ^27^. Of the eight populations in our data that appear to contain mature neurons, four are GABAergic and synapse with glutamatergic neurons. These populations likely include SINs; however, we also identified a glutamatergic population that receives both GABAergic (inhibitory) and glutamatergic (excitatory) input. Thus, populations that demonstrate excitatory-inhibitory connections likely include SINs, but our data suggests these connections occur within the microcircuitry of the tectum as well.

Lastly, we identified a single glycinergic population, OT17, through enrichment of *slc6a9*/GLYT1 which encodes a glutamine transporter responsible for glycine reuptake into the presynaptic cell. GLYT1 may also be expressed by astrocytes serving to regulate neurotransmission, and although previous work found GLYT1 expression in the tectum ^42^, it was unclear if these were neurons or glia. Astrocytes had not been described in zebrafish until 2020, when Chen et al. identified astrocyte-like cells with enrichment of glutamine synthase (*glula*) and glutamate aspartate transporter 1 (*slc1a3b*/GLAST); along with *fgfr3*, and *fgfr4* ^48^, signaling factors required for proper astrocyte growth and function in *Drosophila* ^48,49^. However, OT17 does not appear to highly express any of the genes described by Chen et al (data not shown). Therefore, we believe it likely represents a true glycinergic neuronal population rather than a modulatory astrocyte-like cell.

### Non-neuronal cells of the optic tectum

Not including radial glia, which as previously discussed, may denote immature neuronal populations rather than true mature radial glia; we identified three populations containing non-neuronal cells. Two appear to represent oligodendrocytes, which have been previously identified in the deeper portions of the tectal PVL ^50^. A small population of microglia with an amoeboid morphology has been recently described to localize near neurogenic regions and express the myeloid reporter gene *mpeg1*.*1, as* well as lysosomal genes *ctsla* and *ctsba* ^39^. We found a single small population to express *mpeg1*.*1*, and *ctsba*; as well as *slc7a7*, which functions in microglia development ^51^. From this evidence, we conclude our third non-neuronal cell type to be microglia.

### The optic tectum may provide a useful model for autism pathogenesis studies

A potential link between the SC and autism spectrum disorder (ASD) has been proposed as congenitally blind children are diagnosed with ASD at a higher rate than those born with no vision impairments ^45,52^. As sensory processing impairments typically present in ASD, a structure such as the SC that receives and interprets sensory stimuli poses an interesting avenue for ASD investigations. Utilizing a homologous structure that is highly amenable to genetic manipulations, such as the zebrafish OT, may inform a potential link between the SC and ASD in humans.

Despite clinical heterogeneity and a largely unknown etiology, genetics believed to be a significant contributing factor to ASD. The highly diverse neurexin family of presynaptic cell adhesion molecules is one of the few gene families where every member (NRXN1/NRXN2/NRXN3) is highly implicated in ASD ^53-55^. Diversity can be attributed to alternative promoters generating alpha (long) and beta (short) isoforms, which are then subject to extensive alternative splicing ^56^. Structure is consistent across family members, and alpha/beta isoforms share an intracellular domain as well as one extracellular binding domain ^54^. Although neurexin function is not fully understood, they are hypothesized to interact with postsynaptic neuroligin proteins to induce synapse formation ^56^.

A genome duplication after mammalian and teleost lineages diverged resulted in two zebrafish orthologs for many human genes ^57,58^, including neurexins. Six neurexin genes (*nrxn1a*/*b, nrxn2a*/*b*, and *nrxn3a*/*b)* have been identified but under characterized in zebrafish, although all were shown to be required for synaptogenesis and alpha-*nrxn2a* deficiency was shown to result in increased anxiety ^59-61^. We found the main ortholog of NRXN3 (*nrxn3a*) is differentially expressed within specific populations in the larval OT (Figure 2C-C’’, Figure 6); thus, investigations into a potential SC/OT – ASD link may benefit through the characterizations of neurexins in the OT.

### Transcriptomic characterization of the larval optic tectum facilitates future studies

The amount of information connecting gene expression and tectal cell morphologies is less than ideal; by providing the transcriptomic profiles of tectal cells, we provide the genetic information necessary for future studies to target individual populations and further these associations. This can allow for a variety of experiments such as (1) the use of a population marker to drive reporter gene or calcium indicator expression to observe morphologies or conduct functional assays; or (2) use of previous transgenic lines to drive knock-out or knock-down of marker genes or other highly expressed genes, such as unique transcription factors. Along with providing additional genetic information for future studies, we provide strong evidence that the OT is still undergoing significant developmental fine-tuning. As many functional and molecular studies are typically conducted in relatively young larvae (3 – 10dpf), extending experimental windows to include older larvae/adults may benefit functional OT understanding as a whole. We conclude that although the goal of complete characterization will require still more work, the molecular profiles we have compiled of 25 tectal populations will be a valuable resource in relating transcriptional identity with functional identity.

## ACKNOWLEDGEMENTS

We would like to thank Katie Rondem and Opal Allen at the University of Utah, Huntsman Cancer Institute High Throughput Genomics Core for their sequencing assistance; Sungmin Baek and Tatjana Piotrowski for their cell dissociation and fixation advice; Harold Burgess for providing us with *y304Et*(cfos:Gal4;UAS:Kaede); Roseanne Keeler and Tanya Finken for their excellent care of the BYU Fish Facility; and Daniel Mortensen and Daniel Call in the Biochemistry Department at BYU for their guidance with FAC sorting.

## AUTHOR CONTRIBUTIONS

A.M., A.B., and A.P. conceived and performed experiments. A.S. conceived and provided guidance on experiments. A.M. performed all bioinformatic analyses, and A.M, A.B., A.P., and A.S. wrote the article. J.T.H and B.E.P. provided sequencing and bioinformatic counsel. All authors contributed to article review and revision.

## DECLARATION OF INTERESTS

Authors have no competing interests.

## FUNDING

This work was supported by NICHD:R15HD095737, and internal BYU funding.

## MATERIALS AND METHODS

### RESOURCE AVAILABILITY

#### Lead contact

Further information and requests concerning resources and reagents should be directed to the lead contact, Arminda Suli.

#### Materials availability

There are restrictions on the availability of RNA-FISH probe sets as they are the intellectual property of Molecular Instruments (Los Angeles, CA). We are glad to direct sequence inquiries to their team.

#### Data availability

Raw data generated from this study will be available at NCBI’s Sequence Read Archive (SRA Accession Number: PRJNA779441).

### EXPERIMENTAL MODEL AND SUBJECT DETAILS

#### Zebrafish lines and husbandry

All experiments were performed according to guidelines established by the IACUC review board at Brigham Young University (IACUC Protocol Number: 19-0901). For all experiments, larvae from the *y304Et*(cfos:Gal4;UAS:Kaede) enhancer trap line ^38^ were raised on a 14hr:10hr light:dark cycle and maintained at 28.5°C until 7dpf at which point they were either or harvested for scRNA-seq or fixed in 4% paraformaldehyde preparatory to fluorescent *in situ* hybridization.

### METHOD DETAILS

#### Sample Preparation

15,922 larval zebrafish cells were sequenced from two temporal replicates. Sample preparation across temporal replicates was consistent regarding personnel, equipment, protocol, and sequencing technology.

#### Larvae dissociation and FACS

For each temporal replicate, 7 dpf larvae were anesthetized using tricaine (1:10). Heads were collected via a single cut at the posterior hindbrain and kept on ice for an average of 30 minutes. They were dissociated at 28.5°C in 1mL 1% Trypsin-EDTA, with gentle trituration every 5 minutes for 1 hour. Dissociated cells were filtered with a 70 µm Cup Filcon Filter (BD Biosciences, San Jose, CA, USA) and washed with ice cold DPBS (centrifugation at 5000rpm for 5 minutes at 4°C). Cells were stained with DRAQ5 (1:1000) (biostatus, UK) and DAPI (1:1000) to gate dead cell populations. FAC-sorting was performed at Brigham Young University (Provo, UT, USA) using a FACS Aria Fusion Cell Sorter (BD Biosciences, San Jose, CA, USA). Cells were sorted at a reduced flow rate immediately into methanol to preserve gene expression and pooled together for rehydration and resuspension in preparation for sequencing. Methanol fixed cells were rehydrated via centrifugation in a swinging bucket rotor at 2000g for 5 min at 4°C and resuspended at a concentration of 1000 cells/uL in DPBS with 0.09% BSA.

#### Library preparation and sequencing

Library preparation and sequencing were performed externally at the University of Utah Huntsman Cancer Institute (Salt Lake City, UT, USA) on the 10X Chromium single cell platform (10X Genomics, Pleasanton, CA, USA) using the scRNA-seq 3’ v3.1 Next GEM library preparation pipeline. For each temporal replicate, approximately 30,000 FAC-sorted, methanol-fixed, Kaede^+^ cells were used as input for sequencing; with maximum targeting of 10,000 cells and a sequencing depth of 200M reads.

#### Read alignment and quantification

Raw reads were aligned to version 10 of the zebrafish genome using the 10X Genomics CellRanger (v4.0.0) pipeline. Replicate 1 generated 6,629 barcodes and replicate 2 generated 9,293; each set of barcodes was used to generate a count matrix for downstream analysis (SRA Accession Number: PRJNA779441).

#### Batch Correction and Quality Control

Datasets were considered replicates and not hypothesized to display confounding technical variation. Datasets were examined for batch effects by merging datasets into a single matrix with 15,922 genes, applying a standard analysis workflow, and visualizing the resulting PCA and UMAP plots. A significant amount of overlap between datasets was observed on both PCA and UMAP plots, (SI Figure 2A, 2B), suggesting that merging datasets did not introduce meaningful variation. To further confirm lack of batch effects we determined the expression profile of the canonical habenula marker, *gng8*, and show that habenula cells cluster together across datasets rather than by dataset (SI Figure 2C). For these reasons, batch correction was not deemed necessary, as technical variation between datasets was not sufficient to obscure true biological variation.

Quality control was performed following the workflow outlined by the Harvard Chan Bioinformatics Core in their Single-cell RNA-seq data analysis workshop ^62^. To mitigate the effects of low-quality cells on downstream analyses, cells fulfilling one or more of the following criteria were removed: (1) fewer than 500 UMIs, (2) fewer than 300 genes, or (3) greater than 10% mitochondrial gene content. These parameters identified and removed 1,218 low-quality cells, yielding a final dataset of 14,704 cells. Cell cycle scoring was completed as recommended, and regression of associated genes was not deemed necessary (SI Figure 1) ^62^.

Following quality control, all downstream analyses were implemented using the R package Seurat (v4.0.3) ^63^, following the workflows described on the Satija lab website (https://satijalab.org). Using SCTransform ^64^, data normalization was accomplished by calculating Pearson residuals for the remaining UMI counts via regularized negative binomial regression. SCTransform, a framework for the normalization and variance stabilization of UMI data, has been shown to outperform previous methods by mitigating the effects of technical characteristics such as sequencing depth while preserving biological heterogeneity ^64^. The resulting data matrix was then centered in preparation for dimensional reduction.

#### Dimensional reduction

Pearson residuals were used as direct input for PCA. Forty-three principal components (PCs) were used for dimensional reduction, determined by calculating the percent of variation associated with each PC, and selecting the PC where: (1) 90% of variation was cumulatively explained, and (2) the variation associated with that PC is less than 5%; as recommended by the Harvard Bioinformatics Core ^62^. For visualization purposes, an additional round of dimensional reduction was performed using Uniform Manifold Approximation and Projection (UMAP) post cluster identification.

#### Clustering, marker gene identification, and annotation

Unsupervised clustering was performed with the Seurat function FindClusters(), using the Louvain algorithm with multi-level refinement. Candidate gene markers for each cluster were identified via Seurat’s FindAllMarkers() which implements the MAST test for differential expression. MAST employs a hurdle model tailored to scRNA-seq data by addressing specific characteristics such as stochastic dropout and bimodal expression ^65^. Genes were considered unique markers if they were both highly (>1 log2fc) and widely (>50% of cells) expressed within the cluster of interest compared to all other cells.

Clusters representing the habenula, epiphysis, heart, and olfactory system were identified via known marker genes compiled using the ZFIN database (https://zfin.org/) (SI Figure 3A-B, 4A-B; Table S1). All cells in those clusters were excluded from downstream analyses.

Similarity of OT clusters was calculated using the Seurat function BuildClusterTree(), which constructs a phylogram relating average cells from each cluster based upon a distance matrix constructed in PCA space ^63^.

#### Gene Ontology Analysis

Gene ontology analysis identifies upregulated biological processes through an input list of genes and retrieves the GO terms describing the biological processes(s) each gene is involved in ^66^. Whole data-set gene ontology (GO) analysis was conducted on all significant (p<0.05) differentially expressed genes (DEGs) as identified by the MAST test (Table S2; S3). Note that although the same statistical method was used to determine differentially expressed genes as to nominate candidate cluster markers, the parameters for marker genes were significantly more stringent. For the purposes of GO analysis, a DEG for any given cluster is present in at least 25% of cluster cells and expressed at least .25 log2fc higher in that cluster as opposed to all other cells. GO analysis was not conducted on populations annotated as non-tectal, and any duplicate genes were removed from the final list of DEGs (2,222 genes) which were then used as input. Using PANTHER GO (www.pantherdb.org) ^67^, an online tool maintained by the Gene Ontology Consortium ^68^, we conducted a PANTHER Overrepresentation Analysis for biological process complete annotations using Fisher’s Exact Test with FDR correction ^67^ to identify significant biological processes within the input gene list, providing an effective preliminary tool to explore gene expression data.

#### Gene expression profile analysis

To annotate clusters containing mature neurons, it was reasoned that only functioning, post-mitotic neurons would be actively involved in neurotransmission. Genes involved in various aspects of neurotransmission, such *vamp1* and *sv2a* (vesicle docking/fusion) were manually curated according to their functional designation on ZFIN (https://zfin.org/). For a complete list of neurotransmission processes and their respective genes refer to Table S4. Mature annotation was further confirmed via characterization of neuronal profiles.

To characterize mature and developing neuronal profiles, we performed DEA of genes unique to GABAergic, glutamatergic, dopaminergic, glycinergic, cholinergic, serotonergic, and histaminergic neurons (Table S7). Unique genes for each neurotransmitter were manually curated via ZFIN (https://zfin.org/) and characterized as either presynaptic (describing the neuron itself) or postsynaptic (describing the neuron’s synaptic partners). Presynaptic markers were defined as genes responsible for aspects of neurotransmission such as synthesis, packaging, and presynaptic reuptake, so long as they are unique to a neurotransmitter. To illustrate this process, we demonstrate the characterization of glutamatergic transmission. Unique presynaptic markers include genes such as *slc17a7a/b* (VGLUT1), *slc17a6a*/*b* (VGLUT2), and *slc17a8* (VGLUT3). These genes, present only in glutamatergic neurons, encode vesicular glutamate transporters responsible for loading glutamate into vesicles prior to release at the synaptic cleft. In contrast, *slc1a2a/b* (GLT1) encodes an excitatory amino acid transporter (EAAT) responsible for clearing excess glutamate from the synaptic cleft. However, it cannot be used as a unique presynaptic marker for glutamatergic neurons as glial cells also aid in glutamate removal via GLT1 and other EAATs. Postsynaptic markers for glutamatergic transmission include genes such as *grin1a* and *gria1a*, which build NMDA and AMPA receptors (respectively) present only on the postsynaptic cell membrane.

Oligodendrocyte, microglia and radial glial populations were annotated based upon previously known markers such as *mbpa*/*b, olig1*/*2*, and *mpeg1*.*1* (for a full list of markers see Table S5). Transcription factors were obtained from the UniProt Consortium ^43^ and separated into categories based on review status; only reviewed transcription factors were used in characterization of cell types. Neurogenesis genes were obtained using AmiGO, a web-based software tool ^69^ for browsing the Gene Ontology database ^66,68^ and are available in Table S6. ASD implicated genes were obtained from SFARI Gene, an evolving online database run by the Simons Foundation that evaluates clinical and biological data to score candidate risk genes and CNVs according to their association with ASD ^53^. These scores range from 1 to 3, indicating a gene association of (respectively) high confidence, strong candidate, and suggestive evidence ^53^. For the purpose of differential expression analysis, we curated zebrafish orthologs for all SFARI genes with a score of 1 or 2, excluding those associated with syndromic ASD. This list comprised 316 genes, 298 of which had at least one known zebrafish ortholog; a total of 407 orthologs were used as input for differential expression analysis. All ASD implicated genes with their SFARI scores and orthologs are available in Table S8.

#### RNA velocity

RNA velocity estimates future transcriptional cell states by analyzing the proportion of spliced and unspliced mRNAs of any given gene ^40^. Position sorted BAM files, which contain binary versions of aligned sequence data, were generated by 10X Genomics Cell Ranger 4.0.0 for each replicate ^70^. These BAM files were used as input for velocyto.py (https://velocyto.org/velocyto.py/), which generates a loom file containing spliced and unspliced expression matrices. Each loom file was read using the Seurat function ReadVelocity(), converted into a Seurat object, and merged using the merge() function; the resulting object containing three expression matrices: spliced, unspliced, and ambiguous for all 15,922 cells. The object was then filtered to exclude cells previously removed during quality control of the original merged object.

Following filtration, RNA velocity was computed according to the Satija Lab tutorial for computing RNA velocity on Seurat objects (https://satijalab.org/seurat/), which implements a wrapper around veloctyo.R. Velocity estimations were calculated using the Seurat RunVelocity() function and visualized using the original cluster designations and UMAP embedding obtained during the cluster identification.

#### Hybridization chain reaction RNA-fluorescence in situ hybridization (HCR RNA-FISH)

Kaede^-^ larvae from a *y304Et*(cfos:Gal4;UAS:Kaede;*tyr* ^-/-^) in-cross were grown under standard conditions until 7dpf and fixed in 4% paraformaldehyde (PFA) overnight at 4°C. Following fixation, larvae were washed in PBST (PBS + 10% Tween) for six 15-minute increments and dehydrated for storage in methanol (MeOH) series (5 minutes each: 25% MeOH: 75% PBST; 50% MeOH: 50% PBST; 75% MeOH: 25% PBST; and 100% MeOH). Post dehydration, larvae were transferred into clean 100% MeOH and stored in -20°C. For use, larvae were rehydrated in reverse MeOH series (5 min. each: 100% MeOH; 5% MeOH: 25% PBST; 50% MeOH: 50% PBST; 25% MeOH: 75% PBST, 100% PBST), followed by 3×5min washes in PBST and permeabilization at room temperature in 30ug/mL Proteinase K for 45 minutes. Following rehydration and permeabilization, larvae were post-fixed with 4% PFA for 20 minutes followed by 5×5min washes in PBST to remove the fixative.

HCR split-initiator probe sets were custom designed by Molecular Instruments (https://molecularinstruments.com) using proprietary HCR methodology to detect and fluorescently label target RNA transcripts, while suppressing background automatically ^71^. To maximize targeting of *gng8*, alpha-*nrxn3a*, and *robo4*, probes were designed against shared regions of known variants found on NCBI Gene(https://ncbi.nlm.nih.gov/gene/) ^72^ and the Ensemble database (https://ensemble.org) ^73^. Each probe set consisted of 20 split-initiator probe pairs per target, and all utilized a B1 amplifier, with a 546nm fluorophore label. HCR RNA-FISH experiments were not multiplexed. Probe detection and amplification were performed as recommended by MI protocol for whole mount zebrafish larvae (https://molecularinstruments.com/hcr-rnafish-protocols) with the following optimizations: extended pre-hybridization time (3 hours) and increased probe concentration (16nM). Probe set sequences are considered intellectual property and are proprietary to Molecular Instruments; we direct all sequence inquiries to their technical team.

Following HCR RNA-FISH, larvae were stained with DAPI (1:500) for 3 hours at room temperature, followed by 3×10min washes in PBST. Larvae were then stored in 30% glycerol in PBST away from light at 4°C until imaging. Larvae were embedded dorsal side down in 1.25% low melting agarose on glass bottom culture dishes (MatTek, P35G-0-10C). Images were acquired on an inverted FV1000 Olympus confocal microscope (Olympus Corporation, Japan, Tokyo) with either a 20X or 40XW objective (Olympus, UAPON40XW340) using the 546nm (B1 amplifier) and 400nm (DAPI) lasers. Confocal image data was uploaded into Fiji ^74^ (ImageJ, Bethesda, MA, USA), where images were assigned reference colors and merged for visualization.

### Supplemental Excel Tables

**Table S1. Markers for Non-Tectal Annotation, Related to Figure 1, Figure 2, and STAR Methods**

**Table S2. All Differentially Expressed Genes Identified by MAST Algorithm, Related to Figure 1, Figure 3, and STAR Methods**

**Table S3. Analysis Summary of Gene Ontology PANTHER Overrepresentation Test, Related to Figure 3 and STAR Methods**

**Table S6. Genes Associated with ‘Neurogenesis’ Gene Ontology Term, Related to Figure 3 and Table 1**

**Table S7. Pre- and Postsynaptic Neurotransmitter Markers, Related to Figure 4, Table 1, and STAR Methods**

**Table S8. SFARI Category 1-2 ASD Genes and Reviewed UniProt Transcription Factors, Related to Figure 5, Figure 6, and STAR Methods**

## Notes

### Competing Interest Statement

The authors have declared no competing interest.

### Summary of Updates

This version of the manuscript has been revised to include Sequence Resource Archive (SRA) project reference number.

## References

1. Stein, B. E., & Stanford, T. R. (2013). Chapter 3—Development of the Superior Colliculus/Optic Tectum. In J. L. R. Rubenstein & P. Rakic (Eds.), Neural Circuit Development and Function in the Brain (pp. 41–59). Academic Press. https://doi.org/10.1016/B978-0-12-397267-5.00150-3

2. Krauzlis, R. J., Lovejoy, L. P., & Zénon, A. (2013). Superior colliculus and visual spatial attention. Annual review of neuroscience, 36, 165–182. https://doi.org/10.1146/annurev-neuro-062012-170249

3. Jun, E. J., Bautista, A. R., Nunez, M. D., Allen, D. C., Tak, J. H., Alvarez, E., & Basso, M. A. (2021). Causal role for the primate superior colliculus in the computation of evidence for perceptual decisions. Nature neuroscience, 24(8), 1121–1131. https://doi.org/10.1038/s41593-021-00878-6

4. Basso, M. A., & May, P. J. (2017). Circuits for Action and Cognition: A View from the Superior Colliculus. Annual review of vision science, 3, 197–226. https://doi.org/10.1146/annurev-vision-102016-061234

5. Del Bene, F., & Wyart, C. (2012). Optogenetics: a new enlightenment age for zebrafish neurobiology. Developmental neurobiology, 72(3), 404–414. https://doi.org/10.1002/dneu.20914

6. Vanwalleghem, G., Heap, L. A., & Scott, E. K. (2017). A profile of auditory-responsive neurons in the larval zebrafish brain. The Journal of comparative neurology, 525(14), 3031–3043. https://doi.org/10.1002/cne.24258

7. Gahten, E., Tanger, P., Baier, H. (2005). Visual Prey Capture in Larval Zebrafish Is Controlled by Identified Reticulospinal Neurons Downstream of the Tectum. Journal of Neuroscience, 25(40), 9294-9303. https://doi.org/10.1523/JNEUROSCI.2678-05.2005

8. Nevin, L. M., Taylor, M. R., & Baier, H. (2008). Hardwiring of fine synaptic layers in the zebrafish visual pathway. Neural development, 3, 36. https://doi.org/10.1186/1749-8104-3-36

9. Fero K., Yokogawa T., Burgess H.A. (2011) The Behavioral Repertoire of Larval Zebrafish. In: Kalueff A., Cachat J. (eds) Zebrafish Models in Neurobehavioral Research. Neuromethods, vol 52. Humana Press, Totowa, NJ. https://doi.org/10.1007/978-1-60761-922-2_12

10. Lowe, A. S., Nikolaou, N., Hunter, P. R., Thompson, I. D., & Meyer, M. P. (2013). A systems-based dissection of retinal inputs to the zebrafish tectum reveals different rules for different functional classes during development. The Journal of neuroscience: the official journal of the Society for Neuroscience, 33(35), 13946–13956. https://doi.org/10.1523/JNEUROSCI.1866-13.2013

11. Wong R. O. (1999). Retinal waves and visual system development. Annual review of neuroscience, 22, 29–47. https://doi.org/10.1146/annurev.neuro.22.1.29

12. Nevin, L. M., Robles, E., Baier, H., & Scott, E. K. (2010). Focusing on optic tectum circuitry through the lens of genetics. BMC biology, 8, 126. https://doi.org/10.1186/1741-7007-8-126

13. Heap, L. A., Vanwalleghem, G. C., Thompson, A. W., Favre-Bulle, I., Rubinsztein-Dunlop, H., & Scott, E. K. (2018). Hypothalamic Projections to the Optic Tectum in Larval Zebrafish. Frontiers in neuroanatomy, 11, 135. https://doi.org/10.3389/fnana.2017.00135

14. Robles, E., Filosa, A., & Baier, H. (2013). Precise lamination of retinal axons generates multiple parallel input pathways in the tectum. The Journal of neuroscience : the official journal of the Society for Neuroscience, 33(11), 5027–5039. https://doi.org/10.1523/JNEUROSCI.4990-12.2013

15. Robles, E., Laurell, E., & Baier, H. (2014). The retinal projectome reveals brain-area-specific visual representations generated by ganglion cell diversity. Current biology : CB, 24(18), 2085–2096. https://doi.org/10.1016/j.cub.2014.07.080

16. Xiao, T., Staub, W., Robles, E., Gosse, N. J., Cole, G. J., & Baier, H. (2011). Assembly of lamina-specific neuronal connections by slit bound to type IV collagen. Cell, 146(1), 164–176. https://doi.org/10.1016/j.cell.2011.06.016

17. Meek, J., & Schellart, N. A. (1978). A Golgi study of goldfish optic tectum. The Journal of comparative neurology, 182(1), 89–122. https://doi.org/10.1002/cne.901820107

18. Perry S. F., Ekker M., Farrell A. P., Brauner C. J (2010). Fish Physiology:Zebrafish. Amerstand: Academic Press Elsevier.

19. Yokogawa, T., Hannan, M. C., & Burgess, H. A. (2012). The dorsal raphe modulates sensory responsiveness during arousal in zebrafish. The Journal of neuroscience: the official journal of the Society for Neuroscience, 32(43), 15205–15215. https://doi.org/10.1523/JNEUROSCI.1019-12.2012

20. Filosa, A., Barker, A. J., Dal Maschio, M., & Baier, H. (2016). Feeding State Modulates Behavioral Choice and Processing of Prey Stimuli in the Zebrafish Tectum. Neuron, 90(3), 596–608. https://doi.org/10.1016/j.neuron.2016.03.014

21. Thompson, A. W., Vanwalleghem, G. C., Heap, L. A., & Scott, E. K. (2016). Functional Profiles of Visual-, Auditory-, and Water Flow-Responsive Neurons in the Zebrafish Tectum. Current biology : CB, 26(6), 743–754. https://doi.org/10.1016/j.cub.2016.01.041

22. Scott, E. K., Mason, L., Arrenberg, A. B., Ziv, L., Gosse, N. J., Xiao, T., Chi, N. C., Asakawa, K., Kawakami, K., & Baier, H. (2007). Targeting neural circuitry in zebrafish using GAL4 enhancer trapping. Nature methods, 4(4), 323–326. https://doi.org/10.1038/nmeth1033

23. Scott, E. K., & Baier, H. (2009). The cellular architecture of the larval zebrafish tectum, as revealed by gal4 enhancer trap lines. Frontiers in neural circuits, 3, 13. https://doi.org/10.3389/neuro.04.013.2009

24. Del Bene, F., Wyart, C., Robles, E., Tran, A., Looger, L., Scott, E. K., Isacoff, E. Y., & Baier, H. (2010). Filtering of visual information in the tectum by an identified neural circuit. Science (New York, N.Y.), 330(6004), 669–673. https://doi.org/10.1126/science.1192949

25. Robles, E., Smith, S. J., & Baier, H. (2011). Characterization of Genetically Targeted Neuron Types in the Zebrafish Optic Tectum. Frontiers in Neural Circuits, 5. doi: 10.3389/fncir.2011.00001

26. DeMarco, E., Xu, N., Baier, H., & Robles, E. (2020). Neuron types in the zebrafish optic tectum labeled by an id2b transgene. The Journal of comparative neurology, 528(7), 1173–1188. https://doi.org/10.1002/cne.24815

27. Barker, A. J., Helmbrecht, T. O., Grob, A. A., & Baier, H. (2021). Functional, molecular and morphological heterogeneity of superficial interneurons in the larval zebrafish tectum. The Journal of comparative neurology, 529(9), 2159–2175. https://doi.org/10.1002/cne.25082

28. Kunst, M., Laurell, E., Mokayes, N., Kramer, A., Kubo, F., Fernandes, A. M., Förster, D., Dal Maschio, M., & Baier, H. (2019). A Cellular-Resolution Atlas of the Larval Zebrafish Brain. Neuron, 103(1), 21–38.e5. https://doi.org/10.1016/j.neuron.2019.04.034

29. Farnsworth, D. R., Saunders, L. M., & Miller, A. C. (2020). A single-cell transcriptome atlas for zebrafish development. Developmental biology, 459(2), 100–108. https://doi.org/10.1016/j.ydbio.2019.11.008

30. Niell, C. M., Meyer, M. P., & Smith, S. J. (2004). In vivo imaging of synapse formation on a growing dendritic arbor. Nature neuroscience, 7(3), 254–260. https://doi.org/10.1038/nn1191

31. Niell, C. M., & Smith, S. J. (2005). Functional imaging reveals rapid development of visual response properties in the zebrafish tectum. Neuron, 45(6), 941–951. https://doi.org/10.1016/j.neuron.2005.01.047

32. Pietri, T., Romano, S. A., Pérez-Schuster, V., Boulanger-Weill, J., Candat, V., & Sumbre, G. (2017). The Emergence of the Spatial Structure of Tectal Spontaneous Activity Is Independent of Visual Inputs. Cell reports, 19(5), 939–948. https://doi.org/10.1016/j.celrep.2017.04.015

33. Chen, G., Ning, B., & Shi, T. (2019). Single-Cell RNA-Seq Technologies and Related Computational Data Analysis. Frontiers in genetics, 10, 317. https://doi.org/10.3389/fgene.2019.00317

34. Luecken, M. D., & Theis, F. J. (2019). Current best practices in single-cell RNA-seq analysis: a tutorial. Molecular systems biology, 15(6), e8746. https://doi.org/10.15252/msb.20188746

35. Lush, M. E., Diaz, D. C., Koenecke, N., Baek, S., Boldt, H., St Peter, M. K., Gaitan-Escudero, T., Romero-Carvajal, A., Busch-Nentwich, E. M., Perera, A. G., et al. (2019). scRNA-Seq reveals distinct stem cell populations that drive hair cell regeneration after loss of Fgf and Notch signaling. eLife, 8, e44431. https://doi.org/10.7554/eLife.44431

36. Kölsch, Y., Hahn, J., Sappington, A., Stemmer, M., Fernandes, A. M., Helmbrecht, T. O., Lele, S., Butrus, S., Laurell, E., Arnold-Ammer, I., Shekhar, K., Sanes, J. R., & Baier, H. (2021). Molecular classification of zebrafish retinal ganglion cells links genes to cell types to behavior. Neuron, 109(4), 645–662.e9. https://doi.org/10.1016/j.neuron.2020.12.003

37. Pandey, S., Shekhar, K., Regev, A., & Schier, A. F. (2018). Comprehensive Identification and Spatial Mapping of Habenular Neuronal Types Using Single-Cell RNA Seq. Current Biology : CB, 28(7), 1052–1065.e7. https://doi.org/10.1016/j.cub.2018.02.040

38. Marquart, G. D., Tabor, K. M., Brown, M., Strykowski, J. L., Varshney, G. K., LaFave, M. C., Mueller, T., Burgess, S. M., Higashijima, S., & Burgess, H. A. (2015). A 3D Searchable Database of Transgenic Zebrafish Gal4 and Cre Lines for Functional Neuroanatomy Studies. Frontiers in neural circuits, 9, 78. https://doi.org/10.3389/fncir.2015.00078

39. Silva, N. J., Dorman, L. C., Vainchtein, I. D., Horneck, N. C., & Molofsky, A. V. (2021). In situ and transcriptomic identification of microglia in synapse-rich regions of the developing zebrafish brain. Nature communications, 12(1), 5916. https://doi.org/10.1038/s41467-021-26206-x

40. La Manno, G., Soldatov, R., Zeisel, A., Braun, E., Hochgerner, H., Petukhov, V., Lidschreiber, K., Kastriti, M. E., Lönnerberg, P., Furlan, A., et al. RNA velocity of single cells. Nature 560, 494–498 (2018). https://doi.org/10.1038/s41586-018-0414-6

41. Anadón, R., Rodríguez-Moldes, I., & Adrio, F. (2013). Glycine-immunoreactive neurons in the brain of a shark (Scyliorhinus canicula L.). Journal of comparative neurology, 521(13), 3057–3082. https://doi.org/10.1002/cne.23332

42. Cui, W. W., Low, S. E., Hirata, H., Saint-Amant, L., Geisler, R., Hume, R. I., & Kuwada, J. Y. (2005). The zebrafish shocked gene encodes a glycine transporter and is essential for the function of early neural circuits in the CNS. The Journal of neuroscience: the official journal of the Society for Neuroscience, 25(28), 6610–6620. https://doi.org/10.1523/JNEUROSCI.5009-04.2005

43. UniProt Consortium (2021). UniProt: the universal protein knowledgebase in 2021. Nucleic acids research, 49(D1), D480–D489. https://doi.org/10.1093/nar/gkaa1100

44. Ku, C. C., Wuputra, K., Kato, K., Pan, J. B., Li, C. P., Tsai, M. H., Noguchi, M., Nakamura, Y., Liu, C. J., Chan, T. F., et al. (2021). Deletion of Jdp2 enhances Slc7a11 expression in Atoh-1 positive cerebellum granule cell progenitors in vivo. Stem cell research & therapy, 12(1), 369. https://doi.org/10.1186/s13287-021-02424-4

45. Jure R. (2019). Autism Pathogenesis: The Superior Colliculus. Frontiers in neuroscience, 12, 1029. https://doi.org/10.3389/fnins.2018.01029

46. Boulanger-Weill, J., & Sumbre, G. (2019). Functional Integration of Newborn Neurons in the Zebrafish Optic Tectum. Frontiers in cell and developmental biology, 7, 57. https://doi.org/10.3389/fcell.2019.00057

47. Zheng, W., Geng, A. Q., Li, P. F., Wang, Y., & Yuan, X. B. (2012). Robo4 regulates the radial migration of newborn neurons in developing neocortex. Cerebral cortex (New York, N.Y. : 1991), 22(11), 2587–2601. https://doi.org/10.1093/cercor/bhr330

48. Chen, J., Poskanzer, K. E., Freeman, M. R., & Monk, K. R. (2020). Live-imaging of astrocyte morphogenesis and function in zebrafish neural circuits. Nature neuroscience, 23(10), 1297–1306. https://doi.org/10.1038/s41593-020-0703-x

49. Muñoz-Ballester, C., Umans, R. A., & Robel, S. (2021). Leveraging Zebrafish To Study Bona Fide Astrocytes. Trends in neurosciences, 44(2), 77–79. https://doi.org/10.1016/j.tins.2020.10.013

50. Ito, Y., Tanaka, H., Okamoto, H., & Ohshima, T. (2010). Characterization of neural stem cells and their progeny in the adult zebrafish optic tectum. Developmental biology, 342(1), 26–38. https://doi.org/10.1016/j.ydbio.2010.03.008

51. Rossi, F., Casano, A. M., Henke, K., Richter, K., & Peri, F. (2015). The SLC7A7 Transporter Identifies Microglial Precursors prior to Entry into the Brain. Cell reports, 11(7), 1008–1017. https://doi.org/10.1016/j.celrep.2015.04.028

52. Jure, R., Pogonza, R., & Rapin, I. (2016). Autism Spectrum Disorders (ASD) in Blind Children: Very High Prevalence, Potentially Better Outlook. Journal of autism and developmental disorders, 46(3), 749–759. https://doi.org/10.1007/s10803-015-2612-5

53. Banerjee-Basu, S., & Packer, A. (2010). SFARI Gene: an evolving database for the autism research community. Disease models & mechanisms, 3(3-4), 133–135. https://doi.org/10.1242/dmm.005439

54. Cao, X., & Tabuchi, K. (2017). Functions of synapse adhesion molecules neurexin/neuroligins and neurodevelopmental disorders. Neuroscience Research, 116, 3 9. https://doi.org/10.1016/j.neures.2016.09.005

55. Tromp, A., Mowry, B., & Giacomotto, J. (2021). Neurexins in autism and schizophrenia-a review of patient mutations, mouse models and potential future directions. Molecular psychiatry, 26(3), 747–760. https://doi.org/10.1038/s41380-020-00944-8

56. Ullrich, B., Ushkaryov, Y. A., & Südhof, T. C. (1995). Cartography of neurexins: More than 1000 isoforms generated by alternative splicing and expressed in distinct subsets of neurons. Neuron, 14(3), 497–507. https://doi.org/10.1016/0896-6273(95)90306-2

57. Amores, A., Force, A., Yan, Y. L., Joly, L., Amemiya, C., Fritz, A., Ho, R. K., Langeland, J., Prince, V., Wang, Y. L., Westerfield, M., Ekker, M., & Postlethwait, J. H. (1998). Zebrafish hox clusters and vertebrate genome evolution. Science (New York, N.Y.), 282(5394), 1711–1714. https://doi.org/10.1126/science.282.5394.1711

58. Postlethwait, J., A. Amores, A. Force, and Y. L. Yan, 1998 Chapter 8 The Zebrafish Genome, pp. 149–163 in The Zebrafish: Genetics and Genomics (Methods in Cell Biology, Vol. 60), edited by H. W. Detrich III, M. Westerfield, and L. I. Zon. Academic Press, San Diego, CA.

59. Rissone, A., Monopoli, M., Beltrame, M., Bussolino, F., Cotelli, F., & Arese, M. (2007). Comparative genome analysis of the neurexin gene family in Danio rerio: insights into their functions and evolution. Molecular biology and evolution, 24(1), 236–252. https://doi.org/10.1093/molbev/msl147

60. Shah, A. N., Davey, C. F., Whitebirch, A. C., Miller, A. C., & Moens, C. B. (2015). Rapid reverse genetic screening using CRISPR in zebrafish. Nature Methods, 12(6), 535–540. https://doi.org/10.1038/nmeth.3360

61. Koh, A., Tao, S., Jing Goh, Y., Chaganty, V., See, K., Purushothaman, K., Orbán, L., Mathuru, A. S., Wohland, T., & Winkler, C. (2021). A Neurexin2aa deficiency results in axon pathfinding defects and increased anxiety in zebrafish. Human molecular genetics, 29(23), 3765–3780. https://doi.org/10.1093/hmg/ddaa260

62. Piper, M., Pantano, L., Mistry, M., & Khetani, R. (2020, February 24). Single-cell RNA-seq: Quality Control Analysis. GitHub. https://github.com/hbctraining/scRNAseq/blob/master/lessons/04_SC_quality_control.md.

63. Hao, Y., Hao, S., Andersen-Nissen, E., Mauck, W. M., 3rd, Zheng, S., Butler, A., Lee, M. J., Wilk, A. J., Darby, C., Zager, M., et al. (2021). Integrated analysis of multimodal single-cell data. Cell, 184(13), 3573–3587.e29. https://doi.org/10.1016/j.cell.2021.04.048

64. Hafemeister, C., & Satija, R. (2019). Normalization and variance stabilization of single-cell RNA-seq data using regularized negative binomial regression. Genome biology, 20(1), 296. https://doi.org/10.1186/s13059-019-1874-1

65. Finak, G., McDavid, A., Yajima, M. et al. MAST: a flexible statistical framework for assessing transcriptional changes and characterizing heterogeneity in single-cell RNA sequencing data. Genome Biol 16, 278 (2015). https://doi.org/10.1186/s13059-015-0844-5

66. Ashburner, M., Ball, C. A., Blake, J. A., Botstein, D., Butler, H., Cherry, J. M., Davis, A. P., Dolinski, K., Dwight, S. S., Eppig, J. T., et al. (2000). Gene ontology: tool for the unification of biology. The Gene Ontology Consortium. Nature genetics, 25(1), 25–29. https://doi.org/10.1038/75556

67. Huaiyu Mi, Anushya Muruganujan, Dustin Ebert, Xiaosong Huang, Paul D Thomas, PANTHER version 14: more genomes, a new PANTHER GO-slim and improvements in enrichment analysis tools, Nucleic Acids Research, Volume 47, Issue D1, 08 January 2019, Pages D419–D426, https://doi.org/10.1093/nar/gky1038

68. Gene Ontology Consortium (2021). The Gene Ontology resource: enriching a GOld mine. Nucleic acids research, 49(D1), D325–D334. https://doi.org/10.1093/nar/gkaa1113

69. Carbon, S., Ireland, A., Mungall, C. J., Shu, S., Marshall, B., Lewis, S., AmiGO Hub, & Web Presence Working Group (2009). AmiGO: online access to ontology and annotation data. Bioinformatics (Oxford, England), 25(2), 288–289. https://doi.org/10.1093/bioinformatics/btn615

70. Zheng, G. X., Terry, J. M., Belgrader, P., Ryvkin, P., Bent, Z. W., Wilson, R., Ziraldo, S. B., Wheeler, T. D., McDermott, G. P., Zhu, J., et al. (2017). Massively parallel digital transcriptional profiling of single cells. Nature communications, 8, 14049. https://doi.org/10.1038/ncomms14049

71. Choi, H., Schwarzkopf, M., Fornace, M. E., Acharya, A., Artavanis, G., Stegmaier, J., Cunha, A., & Pierce, N. A. (2018). Third-generation in situ hybridization chain reaction: multiplexed, quantitative, sensitive, versatile, robust. Development (Cambridge, England), 145(12), dev165753. https://doi.org/10.1242/dev.165753

72. Gene [Internet]. Bethesda (MD): National Library of Medicine (US), National Center for Biotechnology Information; 2004 – [cited 2021 Oct 11]. Available from: https://www.ncbi.nlm.nih.gov/gene/

73. Howe, K. L., Achuthan, P., Allen, J., Allen, J., Alvarez-Jarreta, J., Amode, M. R., Armean, I. M., Azov, A. G., Bennett, R., Bhai, J., et al. (2021). Ensembl 2021. Nucleic acids research, 49(D1), D884–D891. https://doi.org/10.1093/nar/gkaa942

74. Schindelin, J., Arganda-Carreras, I., Frise, E., Kaynig, V., Longair, M., Pietzsch, T., Preibisch, S., Rueden, C., Saalfeld, S., Schmid, B., et al. (2012). Fiji: an open-source platform for biological-image analysis. Nature methods, 9(7), 676–682. https://doi.org/10.1038/nmeth.2019

