## Supplemental Figures 1-5;Tables 4-5 for "Single-Cell RNA Sequencing Characterizes the Molecular Heterogeneity of the Larval Zebrafish Optic Tectum"

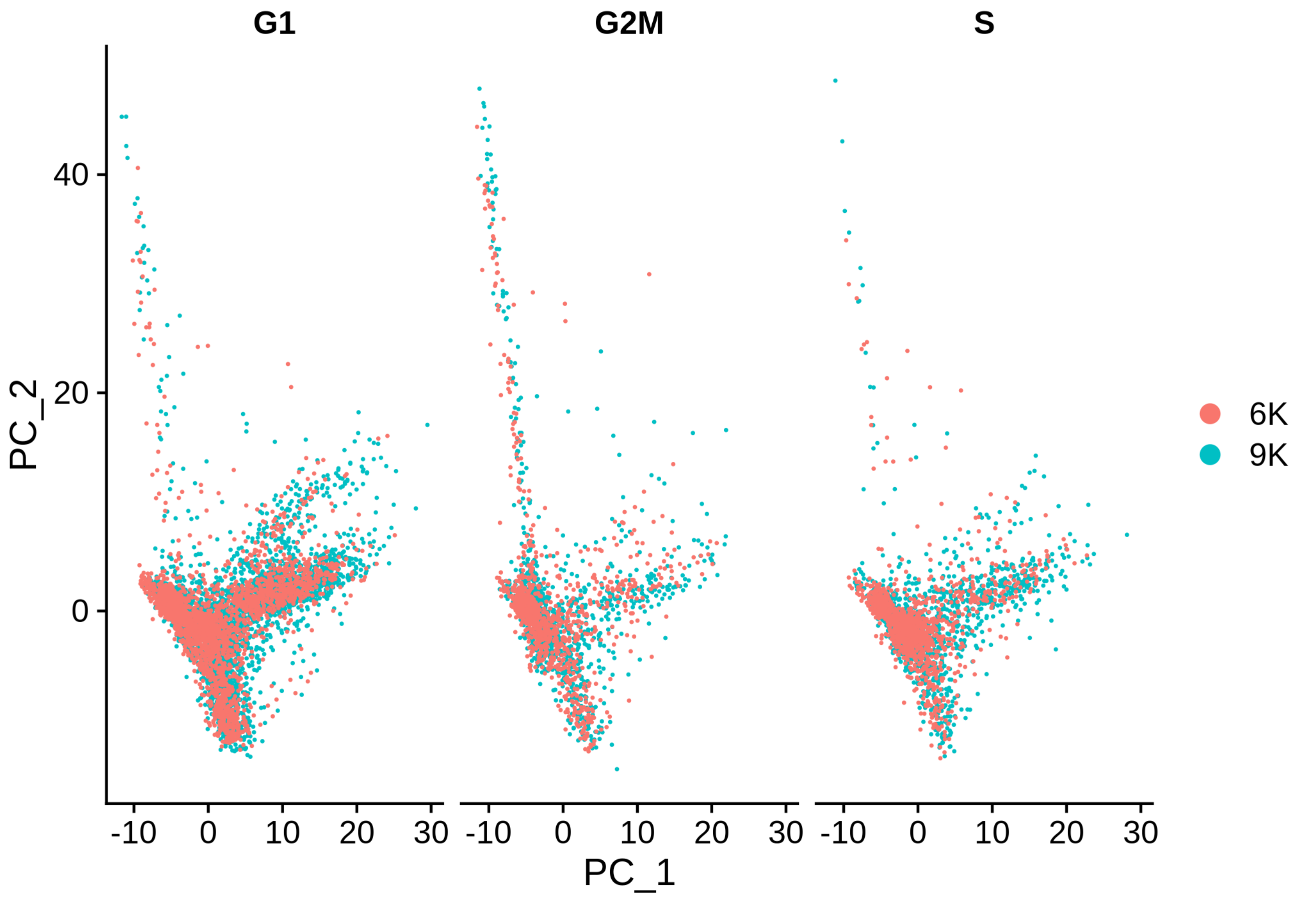


**Figure S1. Quality Control for Cell Cycle Regression, Related to STAR Methods.** Variation in gene expression due to cell cycle can obscure true biological variation. We performed cell cycle scoring for each gene (18222) in the merged dataset to determine if cells group by cell cycle. We found cell cycle did not have an impact on PCA grouping, and regression was not deemed necessary. 6K and 9K denote number of cells sequenced per replicate sequencing replicates

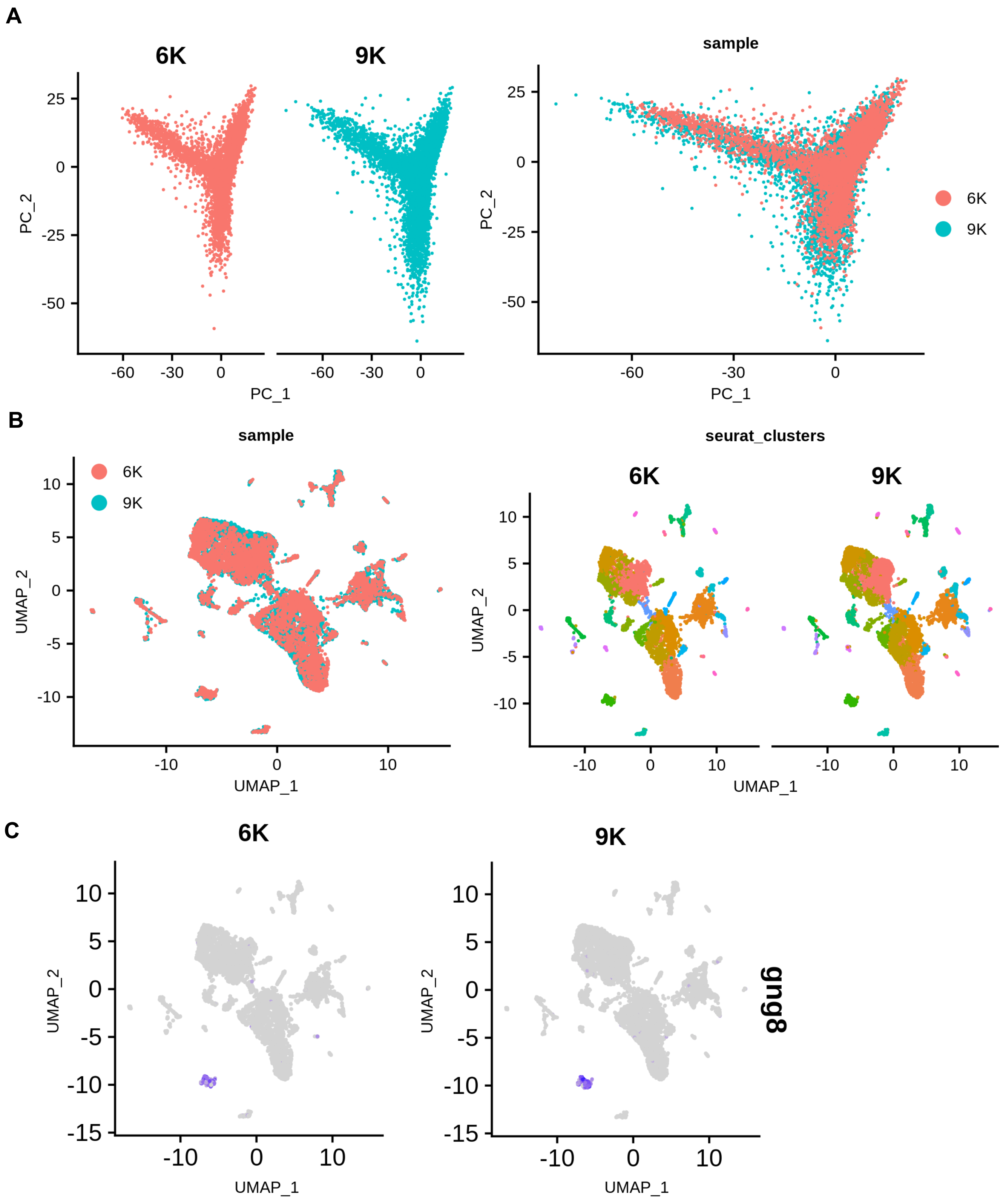


**Figure S2. Quality Control Check for Batch Effect, Related to STAR Methods.** Two temporal replicates of 9,293 and 6,629 cells were used to generate the initial datset of 15,922 total cells; we expect similar variation and do not anticipate batch to obscure meaningful biological variation. A: PCA plots (left:split by replicate, right: overlay) of each replicate show similar grouping, indicating similar variation. B: UMAPs of the merged dataset after clustering (left: sample overlay; right: split). C: UMAPs of the canonical habenula marker gng8 show cells cluster according to cell type rather than replicate, indicating batch effects are not present.

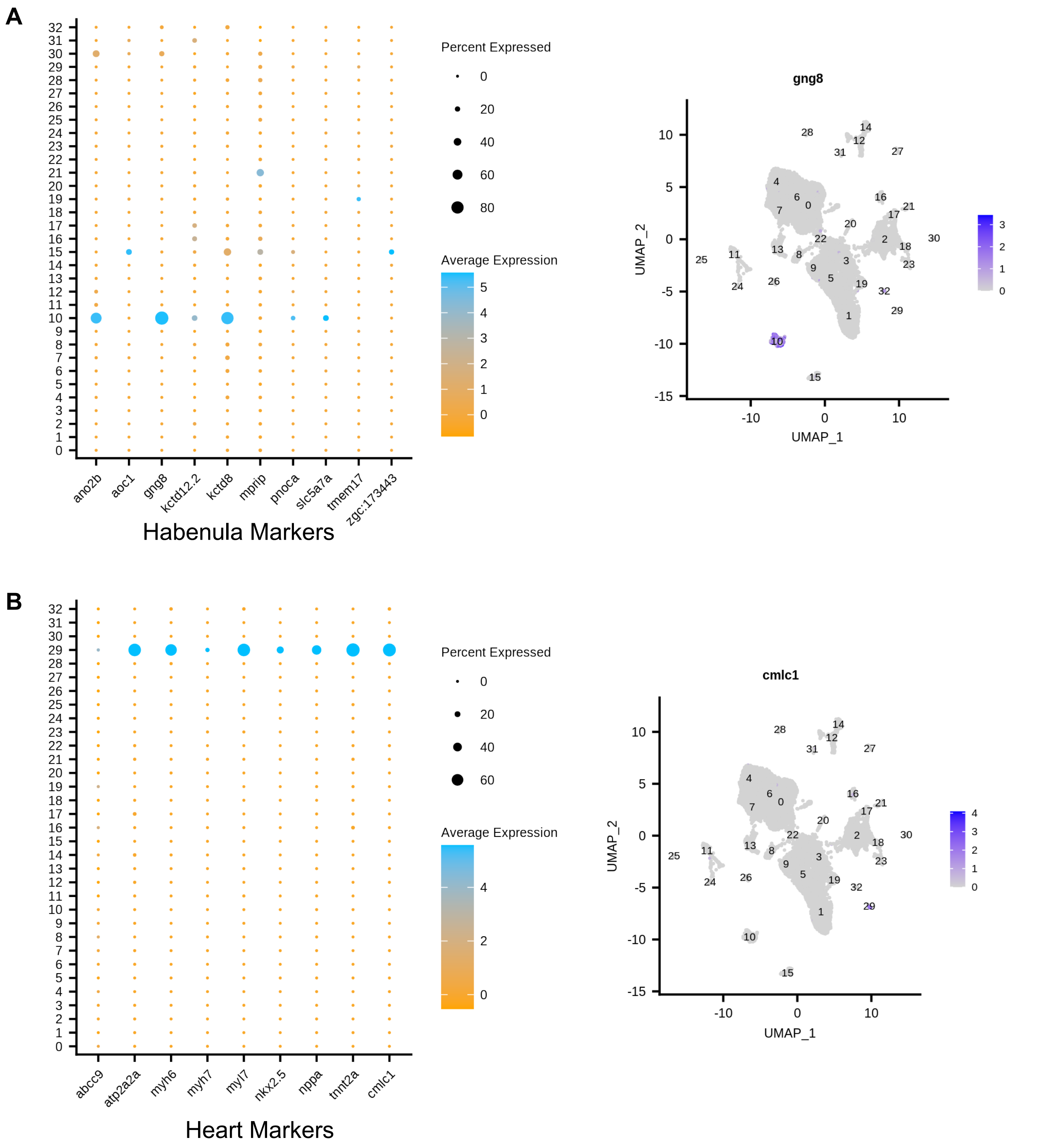


**Figure S3. Annotation of Habenula and Heart Cells, Related to STAR Methods and Figure 1.** To determine putative tectal cells we used known marker genes to exclude populations of non-interest. A: (left) Upregulated expression of habenula genes in cluster 10 nominate it as habenular; (right) expression of *gng8*, canonical habenula marker is restricted to cluster 10. B: Upregulated expression of heart marker genes in cluster 29 nominate it as a heart cluster (left); expression of *cmlc1*, a canonical heart marker, is restricted to cluster 29 (right). See STAR Methods for marker gene curation.

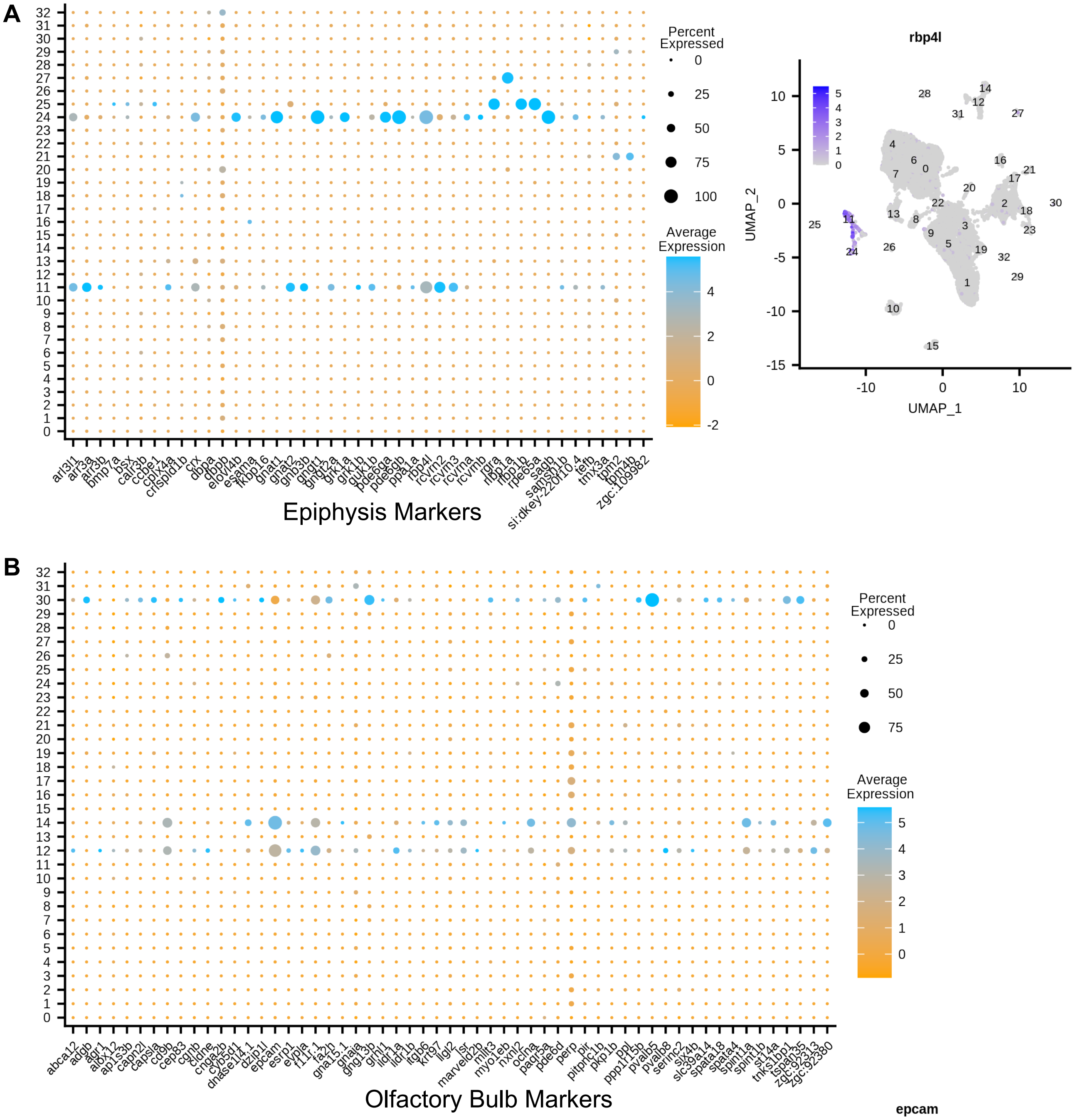


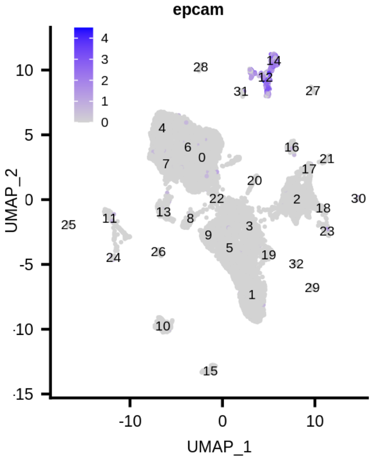


**Figure S4. Annotation of Epiphysis and Olfactory Bulb Cells, Related to STAR Methods and Figure 1.** To determine putative tectal cells, we used known marker genes to exclude populations of non-interest. A: (left) Upregulated expression of epiphysis genes in clusters 11/24/25 nominate them as epiphysal; (right) expression of *rbp4l*, canonical epiphysis/retinal marker is restricted to clusters 11/24/25. B: Upregulated expression of olfactory genes in clusters 12/14/30 nominate them as olfactory (top left); expression of *epcam*, canonical olfactory bulb marker, is restricted to clusters 12/14/30 (bottom right). See STAR Methods for marker gene curation.

Martin et al., 2021

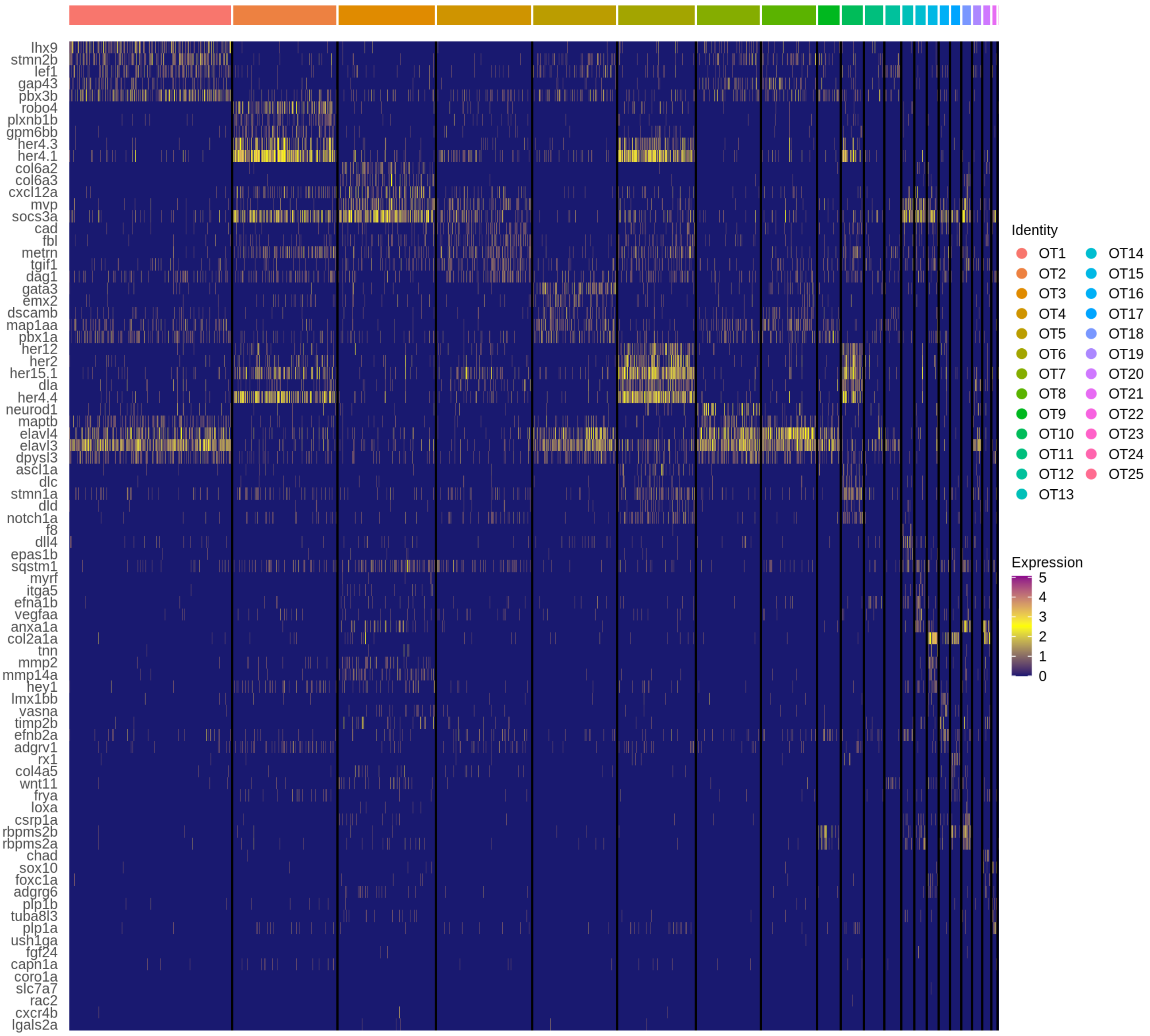


**Figure S5. Top Five Unique Neurogenesis Genes, Related to Figure 3 and Table 1.** Heatmap showing the top five differentially expressed neurogensis genes for each cluster according to log2 fold change; duplicates are allowed but each gene is only represented once. See STAR Methods for differential gene expression parameters.

Table S4. Mature Neuronal Markers, Related to Figure 3, Figure 4, and STAR Methods

| Mature Neurons | |
|---|---|
| Gene | Function |
| sv2a | Predicted to have transmembrane transporter activity. Predicted to be involved in chemical synaptic transmission; neurotransmitter transport; and transmembrane transport. Predicted to localize to cytoplasmic vesicle; integral component of membrane; and synapse. |
| sv2ba | Predicted to have transmembrane transporter activity. Predicted to be involved in chemical synaptic transmission and transmembrane transport. Predicted to localize to integral component of membrane; neuron projection; and synaptic vesicle membrane. |
| sv2bb | Predicted to have transmembrane transporter activity. Predicted to be involved in chemical synaptic transmission; neurotransmitter transport; and transmembrane transport. Predicted to localize to integral component of membrane; neuron projection; and synaptic vesicle membrane. |
| sv2a | Predicted to have transmembrane transporter activity. Predicted to be involved in chemical synaptic transmission; neurotransmitter transport; and transmembrane transport. Predicted to localize to cytoplasmic vesicle; integral component of membrane; and synapse. |
| sv2ca | Predicted to have transmembrane transporter activity. Predicted to be involved in chemical synaptic transmission; neurotransmitter transport; and transmembrane transport. Predicted to localize to integral component of membrane; neuron projection; and synaptic vesicle membrane. |
| vamp1 | Predicted to have SNAP receptor activity and syntaxin-1 binding activity. Predicted to be involved in SNARE complex assembly. Predicted to localize to SNARE complex and plasma membrane. |
| vamp2 | Predicted to have SNAP receptor activity and syntaxin-1 binding activity. Predicted to be involved in SNARE complex assembly. Localizes to cleavage furrow and vesicle. |
| vamp3 | Predicted to have SNAP receptor activity and syntaxin-1 binding activity. Predicted to be involved in SNARE complex assembly. Predicted to localize to SNARE complex and plasma membrane. |
| vamp4 | Predicted to be involved in Golgi ribbon formation; SNARE complex assembly; and microtubule cytoskeleton organization. Predicted to localize to SNARE complex; synaptic vesicle; and trans-Golgi network. |
| vamp5 | Predicted to have SNAP receptor activity and syntaxin-1 binding activity. Predicted to be involved in SNARE complex assembly. Predicted to localize to SNARE complex and plasma membrane. |
| vamp8 | Predicted to have SNAP receptor activity and syntaxin binding activity. Predicted to be involved in SNARE complex assembly and mucus secretion. Predicted to localize to SNARE complex; mucin granule; and plasma membrane. |

Table S5: Glial Markers, Related to Figure 3 and STAR Methods

| Glial Markers | | | |
|---|---|---|---|
| Gene | Link | Expression location | Glial Type |
| her4.1 | http://zfin.org/ZDB-GENE-980526-521#expression | anerior neural rod, ectoderm, nervous system, nerual keep, segmental plate | Radial glia |
| cx43 | http://zfin.org/ZDB-TSCRIPT-090929-1120 | anerior neural rod, ectoderm, nervous system, nerual keep, segmental plate | Radial glia |
| id1 | http://zfin.org/ZDB-GENE-990415-96#summary | brain, ectoderm, germ ring, mesoderm, pleuroperitoneal region | Radial glia |
| s100b | http://zfin.org/ZDB-GENE-040718-290#summary | digestive system, heart, integument, nervous and renal system. | Radial glia |
| fabp7a | http://zfin.org/ZDB-GENE-000627-1#summary | gill, nervous system, eye | Radial glia |
| blbp | http://zfin.org/ZDB-GENE-000627-1#summary | gill, nervous system, eye | Radial glia |
| glula | http://zfin.org/ZDB-GENE-030131-688#summary | cardiovascular, digestive, hematopoietic, muscular, and nervous system | Radial glia |
| si:ch211-251b21.1 | http://zfin.org/ZDB-GENE-060809-5#summary | central nervous system, proliferative region, spinal cord | Radial glia |
| fgfbp3 | http://zfin.org/ZDB-GENE-050208-135#summary | brain, endoderm, hindbrain, midbrain, telencephalon | Radial glia |
| atp1a1b | http://zfin.org/ZDB-GENE-001212-5#summary | central nervous system and nerual tube | Radial glia |
| selenop | http://zfin.org/ZDB-GENE-030311-1#summary | head mesenchyme, pronephric duct, yolk, yolk syncytial layer | Radial glia |
| mdka | http://zfin.org/ZDB-GENE-990621-1#summary | brain, eye, neural tube, neruoectoderm, paraxial mesoderm | Radial glia |
| slc1a2b | http://zfin.org/ZDB-GENE-030131-7779#summary | nervous system, spinal cord, neural tube | Radial glia |
| cd82a | http://zfin.org/ZDB-GENE-030131-2818#summary | central nervous system, otic vesicle, pectoral fin, yolk syncytial layer | Radial glia |
| cxcl12a | http://zfin.org/ZDB-GENE-030318-1#summary | brain, mesoderm, myoseptum, nerual rod, sensory system | Radial glia |
| dhrs12la | http://zfin.org/ZDB-GENE-030131-8104#summary | brain, central nervous system, glial cell, hindbrain | Oligodendrocytes |
| flj13639 | http://zfin.org/ZDB-GENE-030131-8104#summary | brain, central nervous system, glial cell, hindbrain | Oligodendrocytes |
| mbpa | http://zfin.org/ZDB-GENE-030128-2#summary | EVL, nervous system, otic vesicle, periderm | Oligodendrocytes |
| mbpb | http://zfin.org/ZDB-GENE-030429-21#summary | nervous system and polster | Oligodendrocytes |

Table S5: Glial Markers, Related to Figure 3 and STAR Methods

| | | | |
|---|---|---|---|
| mpz | http://zfin.org/ZDB-GENE-010724-4#summary | Rohon-Beard neurons, basal plate midbrain region, central nervous sytem, cranial nerve, oligodendrocytes | Oligodendrocytes |
| mag | http://zfin.org/ZDB-GENE-041217-24#summary | nervous system | Oligodendrocytes |
| olig1 | http://zfin.org/ZDB-GENE-050107-2#summary | cenral nervous system, oligodendrocyte, and trunk | Oligodendrocytes |
| olig2 | http://zfin.org/ZDB-GENE-030131-4013#summary | glioblast, nervous system, neural keel, nerual plate, and neural tube | Oligodendrocytes |
| plp1a | http://zfin.org/ZDB-GENE-001202-1#summary | central nervous system, glial cell, nerual tube | Oligodendrocytes |
| plp1b | http://zfin.org/ZDB-GENE-030710-6#summary | nervous system | Oligodendrocytes |
| swap70b | http://zfin.org/ZDB-GENE-030131-3587#summary | immature eye, mesoderm, nervous system, pectoral fin, and vasculature | Oligodendrocytes |
| sox10 | http://zfin.org/ZDB-GENE-011207-1#summary | glioblast, head, iridoblast, nervous system, nerual crest | Oligodendrocytes |
| zwi | http://zfin.org/ZDB-GENE-030131-8155#summary | nervous system | Oligodendrocytes |
| erbb3a | http://zfin.org/ZDB-GENE-030916-3#summary | epidermis, fin, glial cell, heart | Oligodendrocytes |
| gfap | http://zfin.org/ZDB-GENE-990914-3#summary | anterior neural keel, nervous system, neural tube, neuronal stem cell, and optic vesicle | Oligodendrocytes |
| dhrs12la | http://zfin.org/ZDB-GENE-030131-8104#summary | brain, central nervous system, glial cell, hindbrain | Oligodendrocytes |
| mpeg1.1 | https://zfin.org/ZDB-GENE-030131-7347#summary | myeloid reporter expressed in tectal microglia (cite) | Microglia |
| slc7a7 | https://zfin.org/ZDB-GENE-051127-5#summary | brain microglial cell | Microglia |
| xpr1b | https://zfin.org/ZDB-GENE-060503-266#summary | brain microglial cell | Microglia |
| ctsba | https://zfin.org/ZDB-GENE-040426-2650#summary | lysosomal gene enriched in tectal microglia (cite) | Tectal microglia |
